# Automated 3D reconstruction of the fetal thorax in the standard atlas space from motion-corrupted MRI stacks for 21-36 weeks GA range

**DOI:** 10.1101/2021.09.22.461335

**Authors:** Alena Uus, Irina Grigorescu, Milou P.M. van Poppel, Johannes K. Steinweg, Thomas A. Roberts, Mary A. Rutherford, Joseph V. Hajnal, David F.A. Lloyd, Kuberan Pushparajah, Maria Deprez

**Affiliations:** School of Imaging Sciences & Biomedical Engineering, King’s College London, St. Thomas’ Hospital, London, SE1 7EH, UK; Department of Congenital Heart Disease, Evelina London Children’s Hospital, London, SE1 7EH, UK; Centre for the Developing Brain, King’s College London, London, SE1 7EH, UK

**Keywords:** Fetal MRI, Fetal heart, Automated localisation, Automated pose estimation, Deformable slice-to-volume registration, 41A05, 41A10, 65D05, 65D17

## Abstract

Slice-to-volume registration (SVR) methods allow reconstruction of high-resolution 3D images from multiple motion-corrupted stacks. SVR-based pipelines have been increasingly used for motion correction for T2-weighted fetal MRI since they allow more informed and detailed diagnosis of brain and body anomalies including congenital heart defects (Lloyd et al., 2019). Recently, fully automated rigid SVR reconstruction of the fetal brain in the atlas space was achieved in (Salehi et al., 2019) that used convolutional neural networks (CNNs) for segmentation and pose estimation. However, these CNN-based methods have not yet been applied to the fetal trunk region. Meanwhile, the existing rigid and deformable SVR (DSVR) solutions (Uus et al., 2020) for the fetal trunk region are limited by the requirement of manual input as well the narrow capture range of the classical gradient descent based registration methods that cannot resolve severe fetal motion frequently occurring at the early gestational age (GA). Furthermore, in our experience, the conventional 2D slice-wise CNN-based brain masking solutions are reportedly prone to errors that require manual corrections when applied on a wide range of acquisition protocols or abnormal cases in clinical setting.

In this work, we propose a fully automated pipeline for reconstruction of the fetal thorax region for 21-36 weeks GA range T2-weighted MRI datasets. It includes 3D CNN-based intra-uterine localisation of the fetal trunk and landmark-guided pose estimation steps that allow automated DSVR reconstruction in the standard radiological space irrespective of the fetal trunk position or the regional stack coverage. The additional step for generation of the common template space and rejection of outliers provides the means for automated exclusion of stacks affected by low image quality or extreme motion. The pipeline was evaluated on a series of experiments including fetal MRI datasets and simulated rotation motion. Furthermore, we performed a qualitative assessment of the image reconstruction quality in terms of the definition of vascular structures on 100 early (median 23.14 weeks) and late (median 31.79 weeks) GA group MRI datasets covering 21 to 36 weeks GA range.

## 1. Introduction

Since the emergence of fast acquisition sequences and advanced motion compensation techniques (Malamateniou et al., 2013) MRI has been gradually integrated into clinical practice for imaging of fetal anomalies (Story and Rutherford, 2015; Manganaro et al., 2018).

Single shot turbo spin echo (ssTSE) sequences allow acquisition of each slice in less than a second, which minimises the impact of fetal motion artefacts on image quality. However, inter-slice fetal and maternal motion leads to loss of structural continuity between slices and corruption of 3D volumetric information in 3D stacks.

Slice-to-volume registration (SVR) tools allow reconstruction of high-resolution isotropic 3D images of the fetal brain (Gholipour et al., 2010; Rousseau et al., 2010; Kuklisova-Murgasova et al., 2012) from multiple low-resolution motion corrupted MRI stacks. The more recently proposed deformable SVR (DSVR) method (Uus et al., 2020) designed for correction of non-rigid motion has also been applied for reconstruction of the fetal trunk (Davidson et al., 2021).

Since 2018, rigid SVR (Kuklisova-Murgasova et al., 2012; Kainz et al., 2015) has been employed on regular basis for averaged 3D reconstruction of the 3D fetal heart anatomy as a part of the current clinical practice for diagnosis of fetal congenital heart disease (CHD) (Lloyd et al., 2019, 2021) at Evelina London Children’s Hospital with more than 300 reconstructions performed since 2017. An example of a 30 weeks GA cardiac MRI (CMR) dataset in Fig. 1 shows a set of motion corrupted input stacks (in the through-plane view) and the corresponding SVR-reconstructed 3D fetal thorax which allows detailed segmentation of the heart and examination of fine vascular structures. The reconstructed images are clinically used for detailed 3D diagnosis of congenital anomalies of fetal cardiac vascu-lature, such as suspected coarctation of the aorta and right or double aortic arch (Lloyd et al., 2019).

**Fig. 1.**
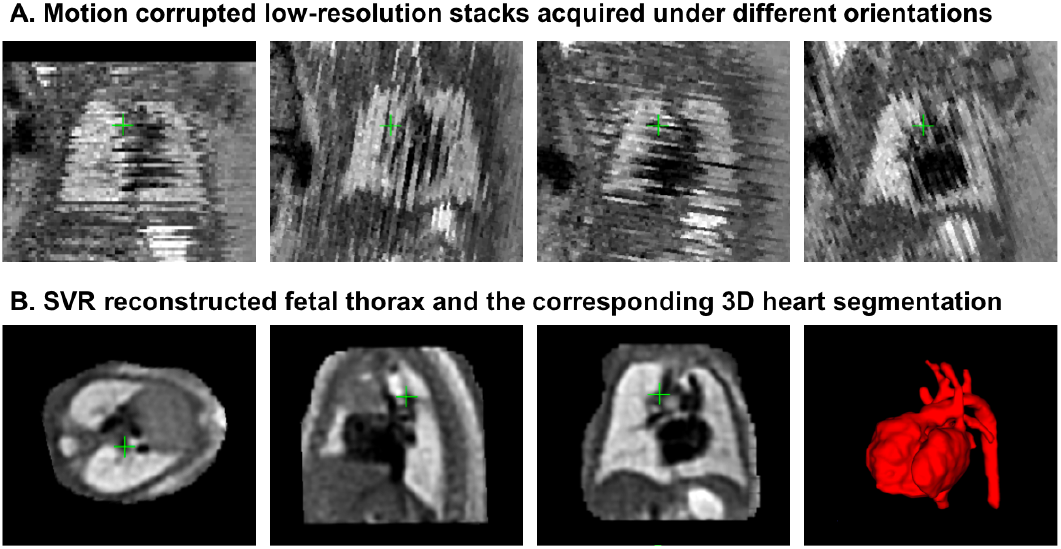
An example of a fetal CMR dataset (30 weeks GA). A: Motion corrupted low resolution stacks acquired under different orientations visualised in the through plane view. B: The corresponding high-resolution SVR-reconstructed fetal thorax and 3D segmentation of the heart and vessels based on the pipeline proposed in (Lloyd et al., 2019).

However, at present, acceptable reconstruction quality can be achieved primarily for the cohort of fetuses from the > 28 weeks gestational age (GA) range. This limitation is caused by the fact that the current SVR methodology (Kuklisova-Murgasova et al., 2012; Kainz et al., 2015) is based on classical registration that cannot resolve large (> 45 - 90°) rotations and translations of the fetal trunk. Early GA cases are particularly prone to large rotations and translations due to the amount of intrauterine space available for manoeuvre (e.g., see Fig. 3.B). For instance, Fig. 2 demonstrates a 23 weeks GA dataset affected by large rotations and translations of the fetus between the stacks which led to failed SVR reconstruction of the thorax.

**Fig. 2.**
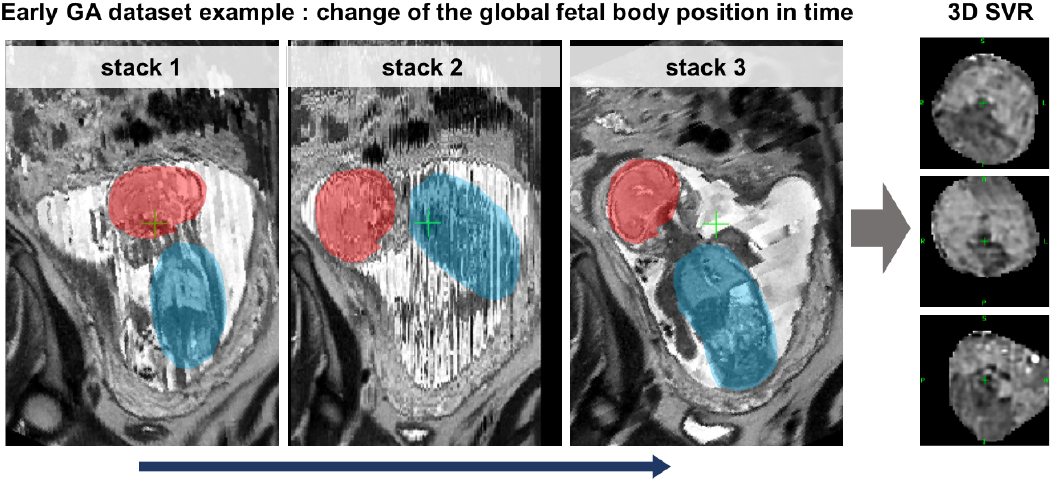
An example of the global change of the fetal trunk (blue) and brain (red) position between stacks during acquisition for an early (23 weeks) GA case. This particular case was affected by severe motion with > 90° rotations and this led to failed SVR reconstruction of the thorax.

**Fig. 3.**
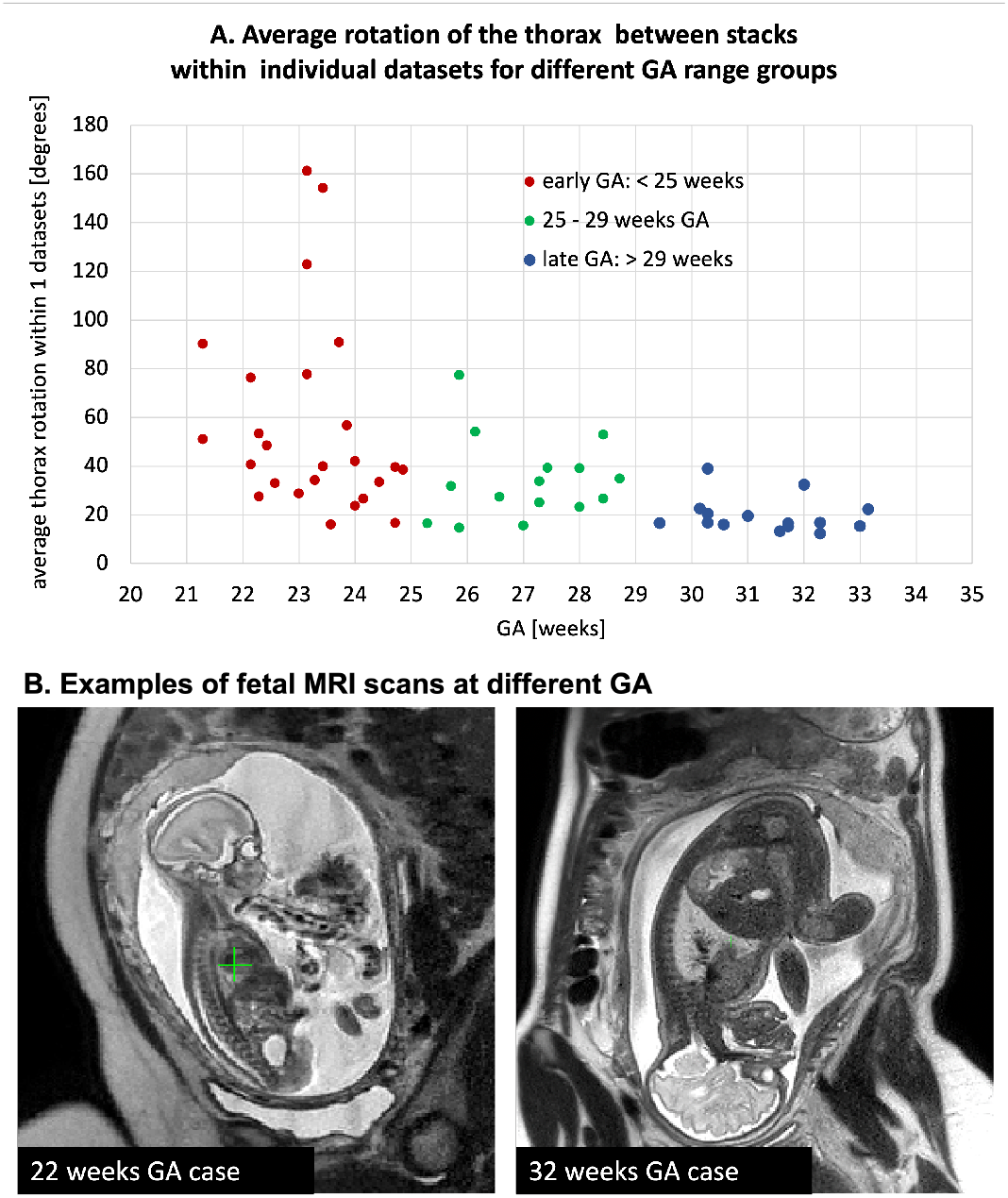
A. Comparison of the degree of the global fetal mobility during MRI acquisition for 55 randomly selected datasets acquired at St. Thomas’s Hospital and Evelina London Children’s Hospital using the same acquisition protocol: average rotation ranges for the fetal thorax ROI (region of interest) between stacks within individual datasets. It includes < 25 weeks GA early (red), 25-29 weeks GA (green), > 29 weeks GA late (blue) groups. B. Examples of fetal MRI scans at 22 and 32 weeks GA.

In general, the degree of motion corruption and its severity varies between datasets. A major proportion of early GA datasets can still be successfully reconstructed using classical SVR or DSVR methods (Uus et al., 2020) if there is a sufficient number of stacks where the fetal trunk is in the same position and only they are selected for the reconstruction. On the other hand, some of late GA cases can also be affected by large rotations due to polyhydramnios when there is too much amniotic fluid around the fetus. The plot in Fig. 3.A shows the average degree of rotation of the fetal thorax position between stacks within individual datasets for randomly selected 40 early and late GA datasets. There is a notable increase in the rotation range for the early GA cases which confirms the limited applicability of the classical SVR-based methods for this cohort.

This limitation was recently addressed for the fetal brain by application of spatial transformer convolutional neural networks (CNN) networks for reorientation of individual 2D slices to the standard radiological atlas space prior to reconstruction (Hou et al., 2018; Salehi et al., 2019) as well as the already reconstructed 3D volumes (Salehi et al., 2019). However, this approach has not yet been applied for motion correction in the fetal trunk ROI. Contrary to the brain, in the fetal trunk ROI, individual 2D slices do not have distinct features required for precise and reliable reorientation to the atlas space. This constitutes a challenge for application of the 2D approaches for pose-estimation of the fetal trunk.

Automation of SVR reconstruction process is another important aspect of general usability and integration into clinical practice. The classical SVR and DSVR methods require manual masks and template stack selection as an input. The existing most efficient solutions for automation of SVR proposed to use 2D CNN slice-wise segmentation for brain masking and intrauterine localisation (Salehi et al., 2018; Ebner et al., 2020). However, in our experience, 2D segmentation often leads to errors due to the insufficient context information or when the object is not present in a stack due to partial coverage. Therefore, the existing automated SVR pipelines reportedly require manual editing and input in a certain proportion of cases. Furthermore, in routine clinical practice, there are also expected inter-site differences in acquisition protocols as well as the coverage of the ROI in input MRI stacks. Full automation without the need for manual inspection of stacks by an operator would require robust localisation.

### 1.1. Related work

During the past decade, different implementations of rigid SVR super-resolution (SR) reconstruction methods were proposed for reconstruction of the fetal brain (Gholipour et al., 2010; Rousseau et al., 2010; Kuklisova-Murgasova et al., 2012; Kainz et al., 2015; Ebner et al., 2020). Based on the approximately rigid motion assumption within the rib cage, SVR was also successfully applied for reconstruction of the averaged 3D fetal heart anatomy (Lloyd et al., 2019). More recently, deformable SVR (Uus et al., 2020) showed to provide improved reconstruction quality for the fetal trunk and placenta ROIs affected by non-rigid motion.

Recently, the limited capture range of the classical registration methods was proposed to be addressed via fetal pose estimation. In this case, the term pose estimation represents computation of the transformation of the object (fetus) the 3D reference world space. The transformation can be retrieved by using either regression or landmark detection approach widely used in computer vision (Wang et al., 2021). The two major existing solutions for the fetal brain pose estimation are based on regression (Hou et al., 2018; Salehi et al., 2019) convolutional neural networks (CNN). They are used for prediction of position of individual 2D slices in the atlas space and the outputs transformations are then used to initialise the SVR reconstruction pipelines. In (Wright et al., 2018), a Long Short-Term Memory (LSTM) network was used for rigid registration of motion-corrected 3D MRI and ultrasound images of the fetal brain and reorientation to the standard space. An alternative landmark-based CNN approach was proposed in (Xu et al., 2019) for pose estimation of the whole fetus based on 15 keypoint landmarks in low-resolution echo-planar imaging (EPI) stacks.

Recently, a series of CNN segmentation-based solutions were proposed for fetal brain localisation and automation of SVR reconstruction. These works employed a 2D UNet (Salehi et al., 2018), a 2D P-net (Ebner et al., 2020) or a 3D V-net for ellipse brain model fitting (Cordero-Grande1 et al., 2019). The 2D output segmentations were then combined into 3D masks and refined using morphological operations and passed to SVR pipelines. In (Li et al., 2020; Fadida-Specktor et al., 2021), a 3D UNet was successfully used for localisation of the whole fetus in EPI and balanced turbo field echo stacks.

### 1.2. Contributions

In this work, we propose a fully automated pipeline for 3D reconstruction of the fetal thorax in the atlas space from motion-corrupted MRI stacks that can capture the full range of fetal motion. It is based on 3D CNN global localisation and landmark-guided pose estimation that allows correction of large rotations and translations that cannot be resolved by the classical registration methods. The additional step for generation of the common template space and rejection of outliers is used in order to account for stacks affected by low image quality or extreme motion. Furthermore, we employ DSVR (Uus et al., 2020) reconstruction rather than rigid SVR used in (Lloyd et al., 2019) since it provides superior performance for the fetal thorax ROI affected by non-rigid motion.

In addition to automation, this solution extends the application of DSVR thorax reconstruction to early GA range cohort that was not previously achievable due to the large rotation motion present in the early GA datasets. The pipeline is evaluated on a series of experiments including both fetal MRI datasets and simulated rotation motion experiments. The general image reconstruction quality with respect to the acceptability for anatomical interpretation is qualitatively evaluated in terms of definition of cardiovascular structures on 100 early and late GA MRI datasets from 21 to 36 weeks GA range.

## 2. Methods

### 2.1. Overview of the algorithm

The proposed pipeline for automated DSVR fetal thorax reconstruction is presented in Fig. 4. In summary, at first, the fetal trunk is globally localised in all stacks using a robust 3D CNN-based segmentation and they are cropped to the trunk ROI. This is followed by segmentation of the thorax, abdomen, heart and liver ROIs and the corresponding centroid landmarks are used for reorientation of all stacks to the standard radiological atlas space. The reoriented stacks are then automatically analysed in terms of similarity and degree of motion corruption in the thorax ROI. Following exclusion of outliers, the template space and the thorax mask are generated as a median average from all preregistered input stacks and masks. The output files are then passed to the standard DSVR reconstruction pipeline (Uus et al., 2020) that produces isotropic high-resolution 3D images.

**Fig. 4.**
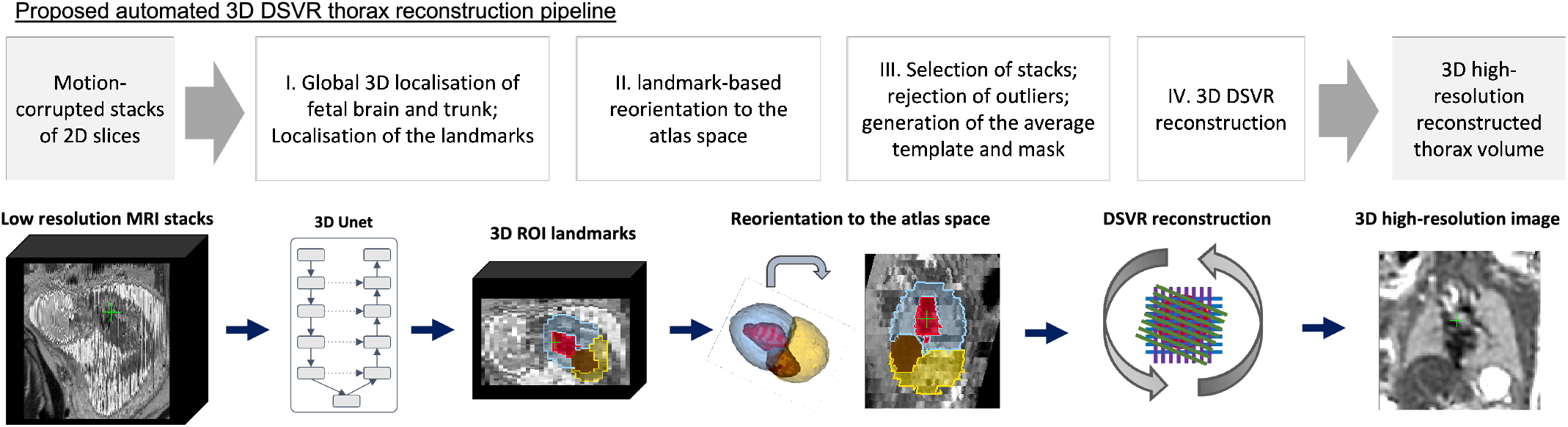
Proposed pipeline for automated DSVR reconstruction of the fetal thorax from motion-corrupted MRI stacks.

### 2.2. Global 3D localisation

In clinical practice, acquired fetal MRI stacks cover different ROIs such as the entire uterus, only the fetal brain or only the trunk. This poses a general challenge to fully automated intra-uterine localisation methods. The existing solutions for localisation of fetal brain in MRI stacks employ 2D slice-wise CNN-based segmentation in combination with morphological operations (Salehi et al., 2018; Ebner et al., 2020). Our experiments showed that 2D UNet localisation of fetal trunk leads to significant errors due to the insufficient context information in individual 2D slices. A comparison of 2D vs. 3D UNet localisation trained on the same (30) datasets is given in Supplemental Figure S1. Furthermore, the slices that contain only the peripheral parts of the fetal brain/trunk and do not have distinctive structure or contrast. We found that the masks generated by the 2D slice-wise approach pipelines are likely to require additional post-processing, manual editing and a certain level of direct quality control from an operator.

In this work, we propose to use a multi-label 3D UNet for simultaneous segmentation-based localisation of the uterus, fetal brain and trunk (Fig. 5) to account for the stacks where only fetal brain or trunk are present and to avoid errors when maternal structures outside the uterus have resemblance to the fetal trunk components. The advantage of 3D multi-component segmentation is the extensive structural information content as well as the mutually exclusive property. In this work, we use only the trunk label for further processing but the brain mask can be potentially used for the whole fetus (head and trunk) reconstruction.

**Fig. 5.**
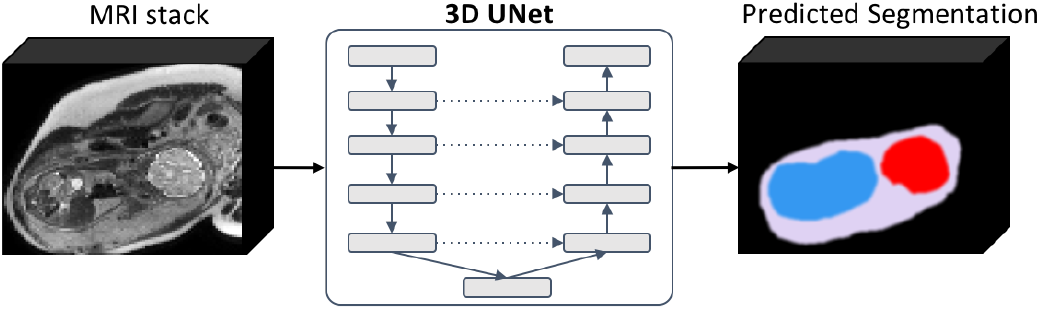
Multi-label 3D UNet network for 3D localisation of the fetal brain (red), fetal trunk (blue) and uterus (lilac) in motion-corrupted 3D MRI stacks.

We employ a classical 3D UNet (Çiçek et al., 2016) architecture with 5 encoding-decoding branches with 32, 64, 128, 256 and 512 channels, respectively. Each encoder block consists of 2 repeated blocks of 3 × 3 × 3 convolutions (with a stride of 1), instance normalisation (Ulyanov et al., 2016) and LeakyReLU activations. The first two down-sampling blocks contains a 2 × 2 × 2 average pooling layers, while the others use 2 × 2 × 2 max pooling layers. The decoder blocks have a similar architecture as the encoder blocks, followed by upsampling layers. This choice of the pooling layers was primarily dictated by the motion corrupted and blurred nature of the input images. The model outputs an *N*-channel 3D image, corresponding to our *N* = 4 classes: background, uterus, fetal brain and trunk. The segmentation network is trained by minimizing a generalised Dice loss (Sudre et al., 2017) using the Adam optimizer with the default parameters (*β*_1_ = 0.9 and *β*_2_ = 0.999), learning rate 0.002 and batch size 2.

As summarised in Fig. 6, following the 3D UNet segmentation step, the trunk labels are extracted with an additional morphological filtering of the largest connected component.

**Fig. 6.**
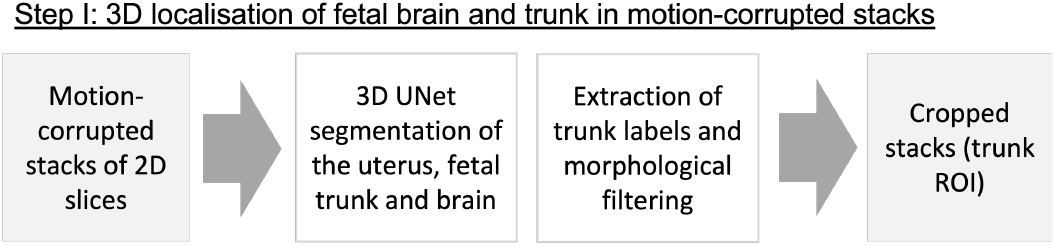
Proposed automated pipeline, Step I: 3D localisation of the uterus, fetal brain and trunk in motion-corrupted stacks.

### 2.3. Landmark-guided pose estimation

As mentioned before, correction of large rotations and translations of the fetal thorax within the same dataset cannot be resolved by the classical rigid registration and poses a particular challenge for processing of the wider GA range MRI datasets. Therefore, integration of the fetal trunk pose estimation step into the pipeline and reorientation of all input stacks to the same reference space is one of the requirements for robust reconstruction performance. In this work, we propose to perform global reorientation of the 3D input stacks rather than 2D slice-wise approach used in (Hou et al., 2018; Salehi et al., 2019). This allows incorporation of 3D spatial information and that minimises the errors for marginal 2D slices with not sufficiently defined structural content.

In summary, the proposed fetal trunk pose estimation step (Fig. 7) is based on automated detection of a set of ROI-specific 3D landmarks (similarly to (Xu et al., 2019)) within the fetal trunk in each stack followed by point-based registration to the atlas space. The output transformations represent the estimated pose of the fetal trunk in the standard 3D space. The 3D landmark-based solution is translation invariant and simultaneously corrects for both rotations and translations.

**Fig. 7.**
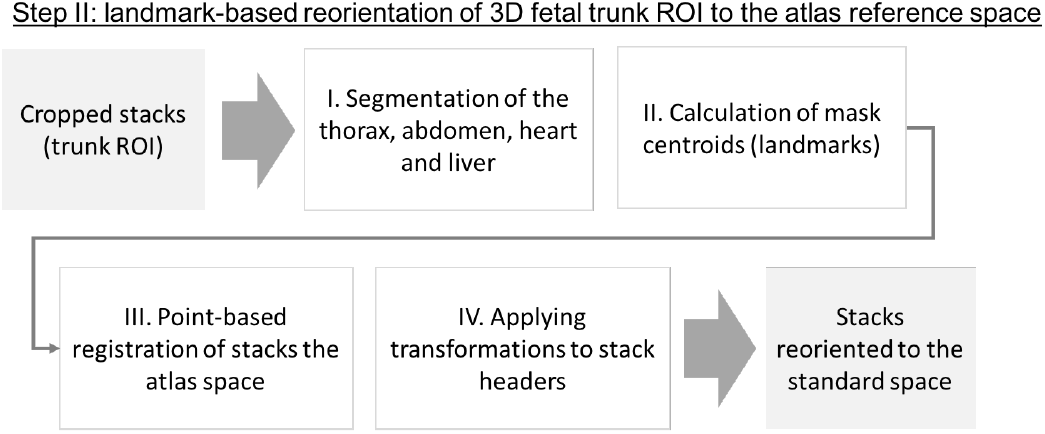
Proposed automated pipeline, Step II: landmark-based 3D fetal thorax pose estimation and reorientation to the atlas space.

We selected centre points of the thorax, abdomen, heart and liver ROI masks as the landmarks. These ROIs are easily identifiable in low-resolution MRI stacks (Fig. 8.A) and are present in both normal and abnormal cases (e.g., missing stomach or lung lesions). While there is an expected inter-subject variability, the position and relative size and position of the selected regions is stable across our target GA.

**Fig. 8.**
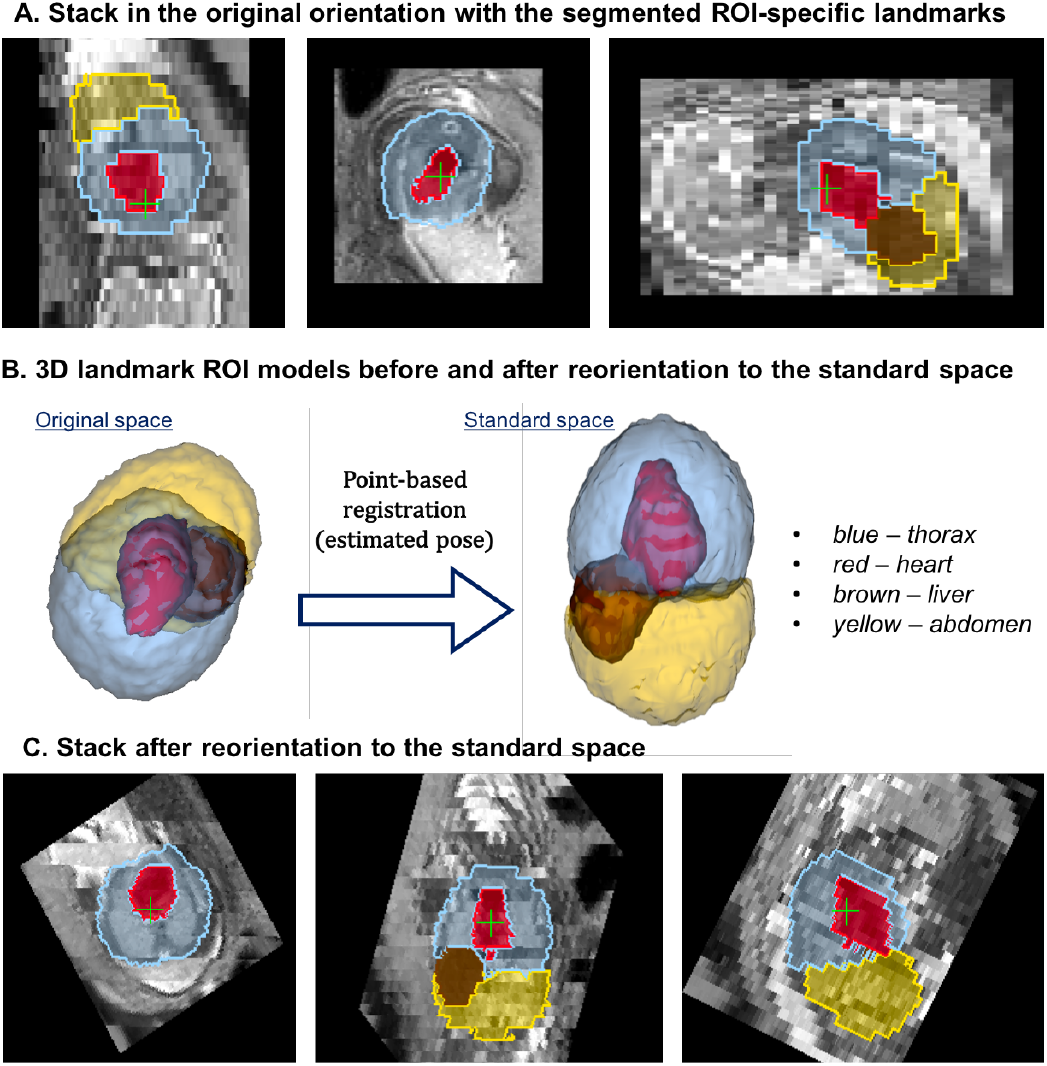
An example of landmark-based reorientation to the atlas reference space: (A) an original motion-corrupted stack in random orientation with detected ROI-specific landmarks, (B) the corresponding 3D models in the original orientation and after transformation to the standard space based on point registration (pose estimation) and (C) the final reoriented stack.

The ROI masks are extracted using the classical 3D UNet (see architecture described in Sec. 2.2) segmentation of the input stacks cropped to the fetal trunk region. The cropping of stacks is required because the inputs to the 3D network are resampled to 128×128×128 grid that leads to loss of resolution for small features for large coverage stacks. This is why we employ hierarchical localisation approach.

The transformations to the standard radiological coordinate system are computed using rigid point-based registration to the same organ centre-point landmarks in the standard radiological atlas space. These transformations represent the estimated fetal pose in every stack.The registration step is performed for every stack and the output rigid transformations are applied directly to the NIfTI header orientation matrices reorienting (estimated pose).

However, taking into account the varying degree of motion corruption and possible inaccuracies in 3D segmentation of the landmarks, the output positions of the landmarks and transformations are not expected to be precise. We also cannot use direct registration to the atlas due to expected inter-subject deviations from the atlas anatomy, especially for abnormal cases. Thus, we introduce an additional rigid registration step which is performed at the next stage of the pipeline.

### 2.4. Automated stack selection and template generation

Selection of the initial template for global registration directly defines the quality of reconstruction outputs (Kuklisova-Murgasova et al., 2012; Uus et al., 2020). Poor template quality (due to either severe motion corruption or a ROI pose different from the majority of stacks) is one of the common reasons for failure of the classical SVR methods. The conventional approach for template selection implies manual inspection of all input stacks by an operator.

At the next step of the proposed automated pipeline, the reoriented stacks are rigidly registered to each other for refinement of the global landmark-based estimated pose transformations and automatically analysed in terms of the mutual similarity and the degree of motion corruption. This is necessary for selection of the most optimal common trunk position and generation of the robust average template and thorax mask required for reconstruction. This step also includes rejection of outlier stacks, which may be affected by misregistration, severe motion corruption, intensity artifacts or the absence of the fetal trunk within the stack coverage.

The pipeline summarised in Fig. 9 includes: (i) refinement of pair-wise stack alignments by rigid registration of all pairs of stacks (the thorax ROI only) initialised by the landmark-based transformations; (ii) calculation of inter-slice motion-corruption (Eq. 1), volume difference (Eq. 3) and mutual stack similarity (Eq. 2) metrics; (iii) selection of the stack with the highest quality and similarity scores 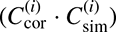 for definition of the common reference space; (iv) reorientation of all stacks to the stack with the highest mutual similarity; (v) rejection of stack outliers based on the computed metrics; (vi) generation of the median template from the selected reoriented stacks; (vii) generation of the average thorax mask from the selected stack masks.

**Fig. 9.**
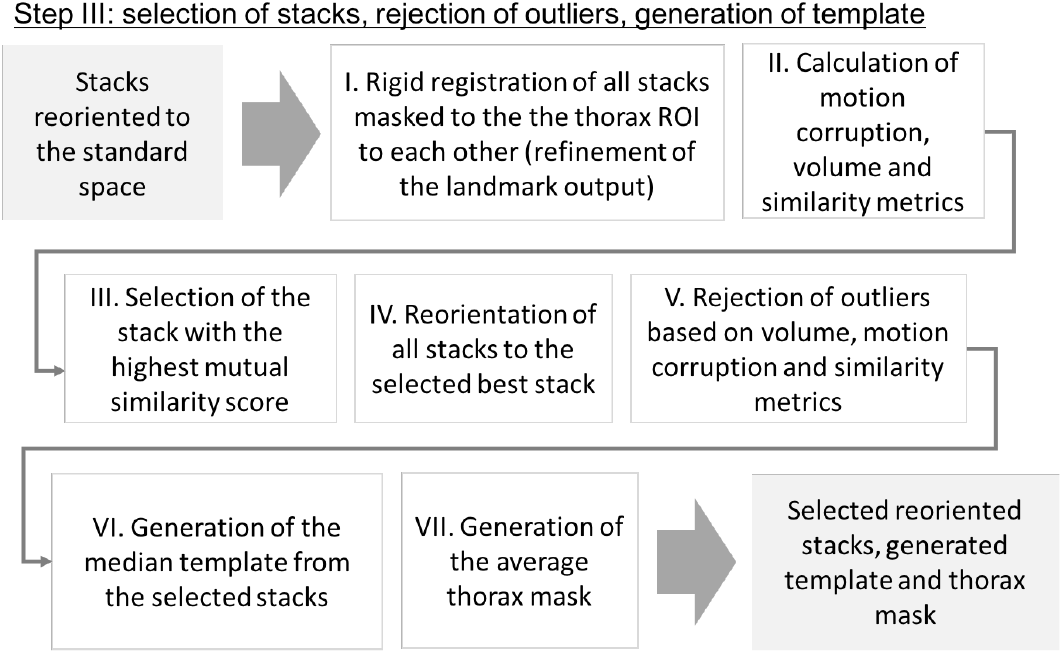
Proposed automated pipeline, Step III: selection of stacks, rejection of outliers and template generation.

**Fig. 10.**
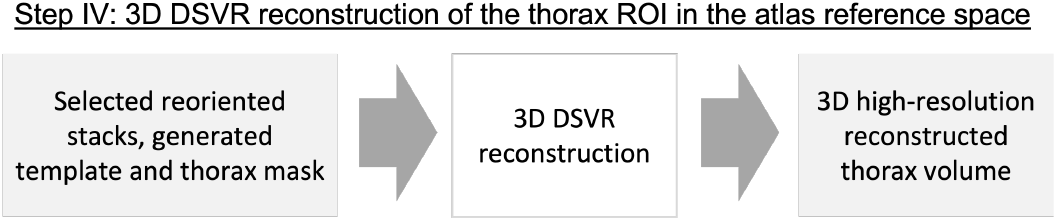
Proposed automated pipeline, Step IV: 3D DSVR reconstruction of the thorax ROI.

For each individual stack (denoted by index *i*, *i* = 1,…, *N*_stacks_) we compute the following metrics: the degree of within-stack motion corruption 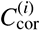, similarity with the rest of the stacks 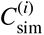 and deviation from the median thorax volume 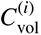. These metrics are computed as:

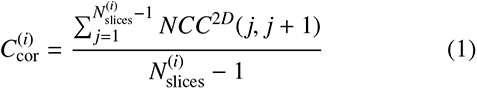

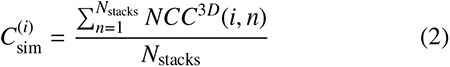

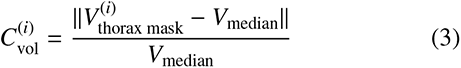

where *NCC* is normalised cross-correlation between sequential 2D slices or individual 3D stacks computed over a non-zero overlapping region, *N*_stacks_ is the number of stacks, 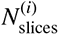 is the number of slices in stack *i* and *V*_median_ and *V*_thorax mask_ are the median and individual stack thorax mask volumes.

The corresponding stack inclusion criteria are as follows:

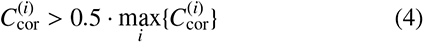

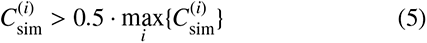

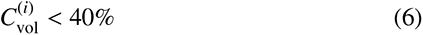

These criteria ensure exclusion of misregistered and severely motion-corrupted stacks as well as stacks with small mask volumes (e.g., when the thorax is absent in the stack).

Following registration of stacks and analysis of the computed metrics, all stacks are reoriented with respect to the stack with the highest mutual stack similarity metric. Following rejection of outliers, the final template and thorax mask are generated as a median average of all remaining stacks.

### 2.5. DSVR reconstruction

As the final step, the selected reoriented stacks along with the generated average template and the mask are passed to the 3D reconstruction step, which is based on our earlier proposed DSVR method. DSVR showed to outperform rigid SVR (Kuklisova-Murgasova et al., 2012) for the fetal trunk ROI affected by non-rigid motion such as bending and stretching. It also includes the structure-based outlier rejection step that minimises the impact of misregistered or low image quality slices on reconstruction results. The output of the reconstruction pipeline is a 3D high-resolution (0.7mm) volume of the thorax ROI in the standard radiological atlas space.

## 3. Implementation

### 3.1. Input data requirements

The proposed DSVR-based reconstruction technique requires a sufficient number of stacks in different orientations, coverage of the fetal head and body ROIs. In the current implementation, the method is operational only for singleton pregnancies.

The ssTSE datasets used in this work have thin slices (2.5mm slice thickness, see Sec.4.1), interleave slice order and negative slice gap to achieve denser oversampling of the ROI. This acquisition protocol was optimised at St.Thomas’ Hospital for imaging and SVR-based reconstruction of both fetal brain and body during the past 5 years and is now used for all clinical and research cases. Even though our acquisitions are performed in this way, the methodology is not restricted to this acquisition protocol. It is independent on the slice gap and acquisition order (ascending or interleaved).

For the protocols with interleaved (e.g., “even-odd”) slice acquisition, input stacks should also be divided into individual “packages” based on the slice acquisition order (e.g., 2 or 4) prior to processing in order to minimise the global change of the 3D fetal position within the stacks. In this case, we divided the stacks into 4 packages with 5 mm slice gap. This step is not required for widely employed clinical thick-slice acquisition format with ascending slice order.

Higher number of stacks and thin slices increase the degree of oversampling of the region of interest and reportedly improves the output image quality for SVR-based methods (Kuklisova-Murgasova et al., 2012; Rubert et al., 2021). In this case, we identified 6 stacks (for the protocol in Sec.4.1) as the minimum for good quality visualisation of vascular structures for both minor and severe motion cases. An illustration of the general impact of the number of stacks on DSVR reconstruction quality is given in Supplemental Figure S2.

The proposed pipeline was designed and trained for black-blood T2-weighted ssTSE datasets optimised for fetal body imaging. And while the performance of DSVR reconstruction is independent on the acquisition parameters, in this work, the localisation networks were trained only on 1.5T datasets with TE=80ms. Although the employed augmentation should potentially partially resolve this limitation, this still might lead to sub-optimal quality due to the differences in tissue contrast.

However, low image quality in terms of SNR levels and severe B1 artefacts is another challenge that affects localisation and reconstruction results and should be addressed separately, either during acquisition or using suitable pre-processing techniques. For the purpose of this work, we exclude datasets with extremely low SNR or severe intensity artifacts.

### 3.2. 3D localisation and pose estimation

#### Software

In summary, the proposed pipeline includes three 3D CNN modules: (i) 4 label 3D UNet for global localisation of fetal trunk and brain (the two other labels are uterus and background); (ii) 3 label 3D UNet for segmentation of the marks global fetal thorax and abdomen landmark (the third label is the background); (iii) 3 label 3D UNet for segmentation of the individual fetal organ landmarks (heart, liver and the third label is the background). The networks (ii) and (iii) were used to define the four landmarks used for reorientation of the stacks. We selected this setup because it was robust for different GAs, in presence of normal functional variation (e.g. variable presence of fluid in the digestive tract, shape of the liver) as well as fetal congenital abnormalities (such as diaphragmatic hernia or renal anomalies).

In our implementation, we used two separate networks to segmentation of the landmarks, one to segment thorax and abdomen, and the other for heart and liver. In order to account for overlapping of the outputs, the heart and liver labels are subtracted from the thorax and abdomen labels. This approach produced very similar results to a single network with 4 landmarks.

The networks were implemented in PyTorch^2^ based on the classical 3D UNet architecture (Çiçek et al., 2016) with TorchIO (Perez-García et al., 2020) augmentation. The employd augmentations included: bias field, 360° rotations, motion artifacts. The selected 128 × 128 × 128 grid size due to the varying ROI coverage in stacks and the size of the fetus with respect maternal structures. All input stacks are resampled and padded to this grid size prior to processing. The code is available online at SVRTK Fetal MRI Segmentation repository^3^.

#### Training the network for global localisation of the fetus

For the global 3D localisation step, 32 fetal CMR CHD MRI datasets from the 28-32 GA range were used for training (318 stacks) and 3 for validation (36 stacks). The uterus, brain and trunk masks were created manually for one of the stacks within each dataset and then propagated to the rest using rigid registration. All resulting masks were visually inspected and corrected, when required. The training was performed for 800 epochs with TorchIO augmentation (textcolorbluerandom with 0.3 probability: bias field, 360^°^ rotations, noise, motion artifacts).

#### Training thorax and landmark segmentation networks

For the thorax, abdomen and organ (heart and liver) ROI segmentation, 65 fetal MRI datasets with normal anatomy from 25 to 32 GA range were used for training and 5 for validation. Rather than using the original motion-corrupted stacks, we used 3D DSVR reconstructed fetal trunk ROI images obtained from a subset of cases from the outputs of (Uus et al., 2020) analysis. The masks were created using label propagation from a generated average fetal trunk atlas followed by manual refinement, if required. The training of the networks was performed for 500 epochs each with TorchIO augmentation (random with 0.3 probability: bias field, 360^°^ rotations, anisotropy, noise, motion artifacts).

### 3.3. Reorientation, stack selection and DSVR reconstruction

The landmark-based reorientation is based on the classical 3D point rigid registration method (Arun et al., 1987) implemented using MIRTK library ^4^. The reference point landmarks were defined in the average fetal trunk atlas reoriented to the standard radiological space.

The step for automated selection of the stacks for reconstruction and generation of the template was also implemented based on MIRTK library and is available as a part of SVRTK package^5^ as *stackselection* function.

### 3.4. Software and hardware requirements

The full compiled pipeline will be available at the SVRTK docker repository ^6^ (*fetal_thorax* tag) after publication of the article.

The recommended hardware configuration is 16 GB GPU, 32-64 GB RAM and 6-12 CPU cores. The total processing time varies between 20 and 60 minutes depending on the ROI size (defined by GA), number of stacks, input and output resolution and system configuration.

## 4. Experiments and results

### 4.1. Fetal MRI data

The fetal MRI data used in this work include 85 datasets acquired under the iFIND^7^ project at St. Thomas’s Hospital, London [REC: 14/LO/1806] and 93 datasets acquired as a part of the clinical fetal CHD CMR service at Evelina London Children’s Hospital [REC: 07/H0707/105]. The datasets were collected subject to the informed consent of the participants. The inclusion criteria for the datasets were: singleton pregnancy, no extreme SNR loss and ≥ 6 input stacks.

The acquisitions were performed on a Philips Ingenia 1.5T MRI system using the same protocol without any maternal sedation: ssTSE sequence with TR= 15000ms, TE=80ms, voxel size 1.25 × 1.25 × 2.5mm, slice thickness 2.5mm, slice spacing 1.25mm and interleaved slice order. The stacks were acquired under different orientations and different ROI coverage, with 100-160 slices per stack, depending on GA and orientation. Each of the datasets contains 6-13 T2-weighted stacks with minimum 4 different orientations and covering different ROIs (whole uterus, brain or trunk only). This acquisition protocol was optimised at St.Thomas’ Hospital for both conventional 2D fetal imaging and SVR-based reconstruction over the past 5 years and is now used for all clinical and research cases. The higher number of sliced (negative gap) increases the degree of oversampling of the region of interest thus improving definition of small features.

The experiments in Sections 4.2-4.6 were based on different setups depending on the availability of the segmentations for training. An additional description of how the datasets were used for different parts of the quantitative and qualitative assessment steps is given Supplemental Table ST1. For each of the experiments we used separate datasets for training, validation and testing. The datasets used for quantitative evaluation were not used in training or validation of the networks.

### 4.2. Automated global 3D localisation: fetal MRI datasets

The first step of the pipeline (Sec. 2.2, CNN module 1) for global localisation of the fetal trunk in raw stacks was evaluated on 16 fetal MRI datasets randomly selected from the early (8 cases, ¡ 24 weeks) and late (8 cases, 28-33 weeks) GA cohorts. Each of the datasets contains 9 - 13 stacks (with 164 stacks in total) acquired under minimum 7 different orientations and with different ROI coverage (uterus, brain+trunk or brain or trunk only).

We compared localisation performance of the 3D multi- (*uterus* + *trunk* + *brain*) and single- (*trunkonly*) label 3D UNet cases respect to the fetal trunk label.

For 8 late GA datasets for which we had ground truth labels, the quantitative evaluation (for the fetal trunk) included: the centroid distance (*d*[*mm*]), the false positive rate (*FPR*) and the manually graded localisation quality scores (*LQS*) defined as {*0-incorrect; 1-partially correct; 2-correct*}, see Tab.1. The high centroid distances and FPR values in the baseline 3D UNet results correspond to wrong localisation outputs, while differences in small values are not informative, because the ground truth (GT) masks are not precise and are affected by motion-corruption. For the 8 early GA datasets, we performed LQS evaluation. In this experiment, the uterus and brain labels are not used for evaluation purposes but only to improve localisation by excluding external structures.

**Table 1.** Evaluation of the fetal trunk localisation quality of the multi-label 3D UNet in comparison to a single-label 3D UNet. The results are statistically significant with *p* < 0.01.

| Metric | 3D UNet (3 labels) | 3D UNet (1 label) |
| --- | --- | --- |
| **28-32 weeks GA datasets** |  |  |
| $d[mm]$ | $6.75 \pm 4.68$ | $18.17 \pm 26.38$ |
| $FPR \times 10^3$ | $0.154 \pm 0.192$ | $0.537 \pm 0.944$ |
| LQS | $1.91 \pm 0.33$ | $1.64 \pm 0.75$ |
| **< 24 weeks GA datasets** |  |  |
| LQS | $1.88 \pm 0.27$ | $1.50 \pm 0.70$ |

The multi-label 3D UNet correctly localised the trunk in all datasets (and nothing was detected in brain-only stacks where the trunk was absent) and provided high localisation quality scores. For late GA datasets, fetal trunk localisation using the single label scenario failed in 16% of the cases, primarily in the stacks where only the brain was present. For early GA datasets, the localisation scores demonstrated that 3-label solution also improved the performance and the trunk was correctly localised with only minor segmentation inconsistencies (1.5 scores for slightly larger mask). For the single label case, the trunk was incorrectly localised in the majority of the brain only stacks (22%) leading to lower LQS.

This confirms the feasibility of the proposed multi-label 3D segmentation approach for automated SVR reconstruction pipelines.

An example of the 3D trunk localisation results for two investigated scenarios is shown in Fig. 11 for three stacks with different ROI coverage. Both multiple- and single-label networks successfully localised the thorax in the whole uterus and trunk-only stacks but the single-label 3D UNet failed in the case of the brain only coverage.

**Fig. 11.**
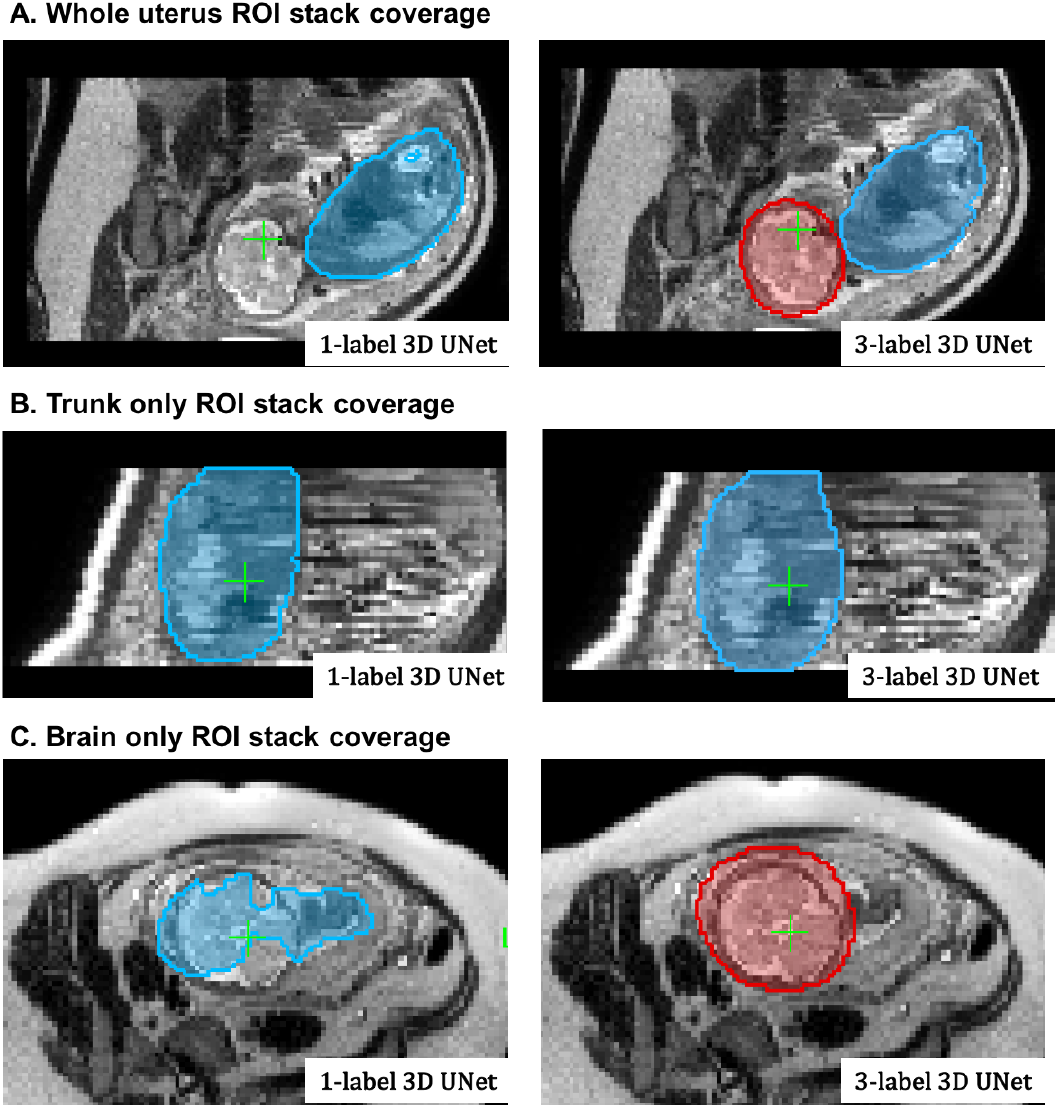
An example of localisation results using multiple- (3) and singlelabel 3D UNet in three stacks with different ROI coverage: the whole uterus (A), trunk only (B), brain only (C). The segmentation outputs are visualised as blue (trunk) and red (brain) overlays.

### 4.3. Automated 3D pose estimation: localisation of landmarks

As a part of the pose estimation pipeline (Sec. 2.3), we performed quantitative assessment of organ landmark localisation on 50 randomly selected stacks covering fetal trunk from 50 different early and late GA datasets originally used for qualitative evaluation (see Fig. 17.B). Before analysis, the stacks were cropped to the trunk ROI based on the output of global localisation.

The reference ground truth labels (thorax, heart, stomach and liver) were created using atlas-based label propagation followed by manual refinement, if required. Comparison with the outputs from the networks (Ses. 3.2) shown in Fig. 12.A-D was performed based on Dice, sensitivity, specificity, centre-point distance and localisation quality score.

**Fig. 12.**
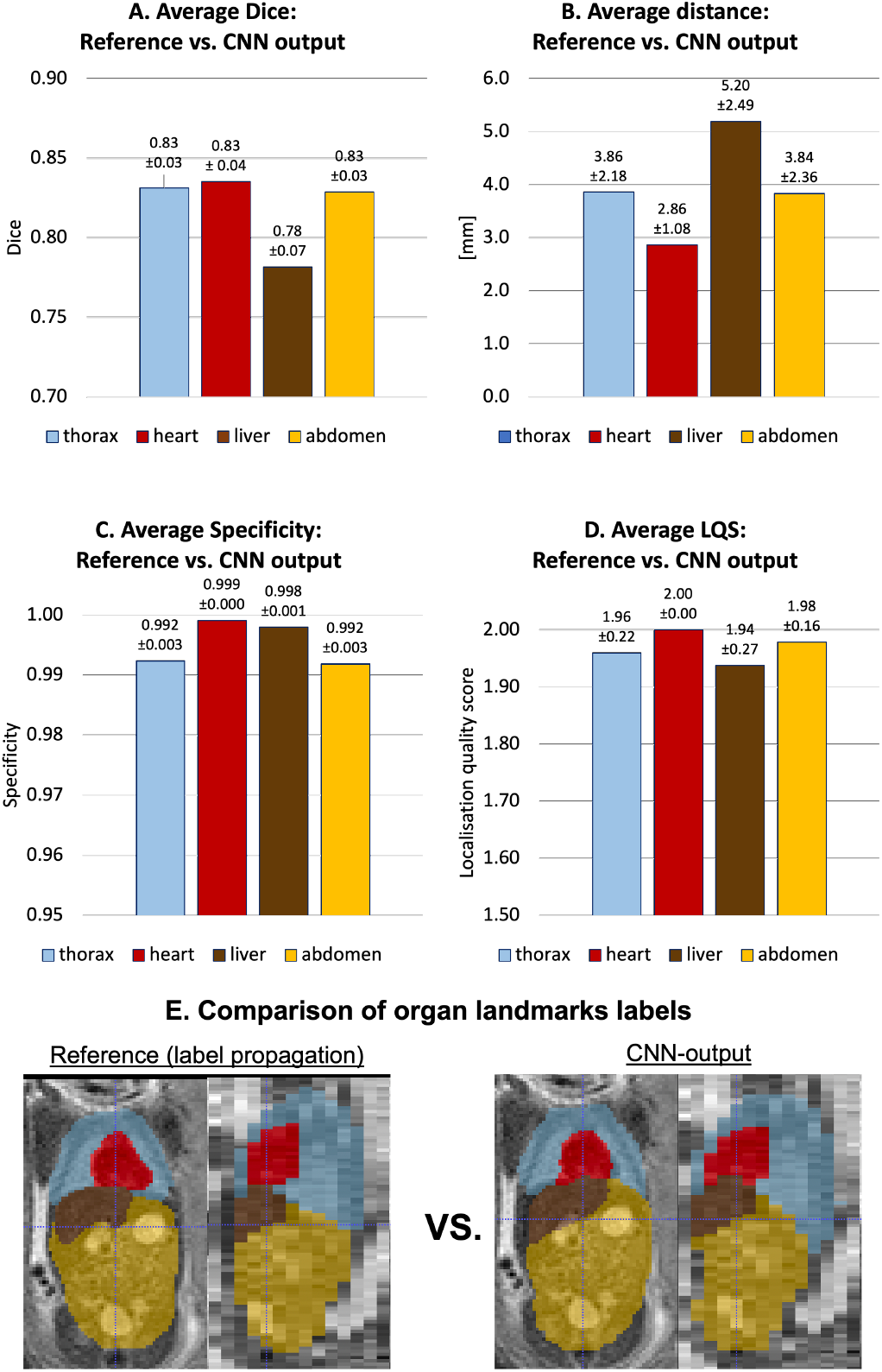
The results of organ localisation for 50 stacks from early and late GA cohort (A-D). An example of reference (based on atlas label propagation) vs. network output for one of the early GA stacks is shown in (E).

The relatively average Dice results (varying between 0.75 and 0.85) are related to the fact that the imperfection of the original input labels used for training and motion-corruption of the input stacks. However, the high specificity and low centre-point distance values confirm the high localisation quality scores. Visual assessment of results (LQS) showed that localisation successful for all stacks and all labels with minor errors (additional) affecting thorax, liver and abdomen. A the same time it did not produce extreme deviations in the compute centroid distance with the maximum ~ 12*mm* error for the liver label (primarily due to shape variability and similarity to the spleen), which is still within the acceptable range for reorientation.

### 4.4. Automated 3D pose estimation: simulated experiment

In order to assess the capture range of the proposed pose estimation approach (Sec. 2.3), we simulated rotations from whole 360 degrees range on 5 normal anatomy datasets from 29 - 32 week GA range without significant motion corruption. Each dataset contains 7 stacks without significant SNR loss or intensity and motion artifacts. All stack were cropped to the dilated trunk mask and globally reoriented to the atlas space with the origin set to zero.

At first, for each of these datasets, one of the stacks was selected as a template and the remaining were additionally registered to it to ensure that the thorax is exactly in the same position.

Next, the rotation motion was simulated by rotating six of the remaining stacks in X, Y and Z direction with the same ± angle. The following rotation angles were selected: {0; 15; 30; 45; 60; 75; 90; 105; 120; 180} degrees in order cover the whole range and identify the limit of the classical rigid registration. An example of simulated rotations is given in Fig. 13.

**Fig. 13.**
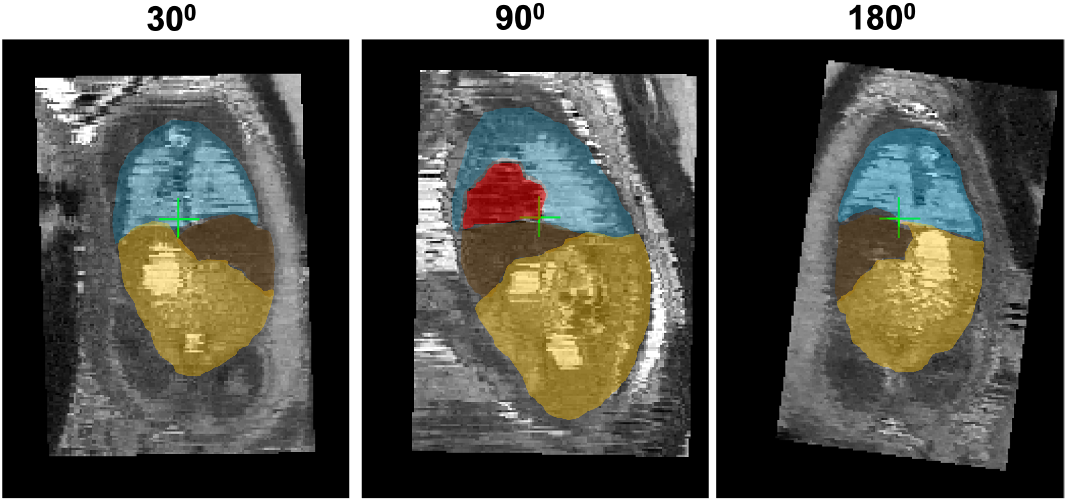
An example of simulated 30, 90 and 180 degrees rotations along one axis applied to a stack cropped to the fetal trunk ROI. The displayed landmark labels include: thorax (blue), heart (red), liver (brown) and abdomen (yellow).

Next, for each of the datasets, we run registration of the rotated stacks to the template using the following three approaches: (i) the classical gradient descent based rigid registration (from MIRTK package), (ii) the proposed automated landmark-based pose estimation approach (Sec. 2.3) and (iii) the combination of the previous two methods with the landmark-based output used for initialisation of the classical rigid registration.

The graphs in Fig. 14 show the average global 3D NCC between the template stack and transformed registered stacks in the masked thorax ROI, calculated over all stacks in all datasets. The drop in NCC values to almost zero for > 75 degrees rotations confirms the limited capture range of the classical rigid registration method (red, average NCC = 0.243 ± 0.207). The consistent intermediate level of similarity for the purely landmark-based output (blue, average NCC = 0.177 ± 0.005) with small standard deviation confirms that this method is rotation-invariant and provides approximate global reorientation to the atlas space. The fact that the NCC values are lower than the classical registration outputs for < 90 degree range indicates that the alignment is not very accurate. This is caused by differences in centre-point positions in segmented 3D landmark ROIs.

**Fig. 14.**
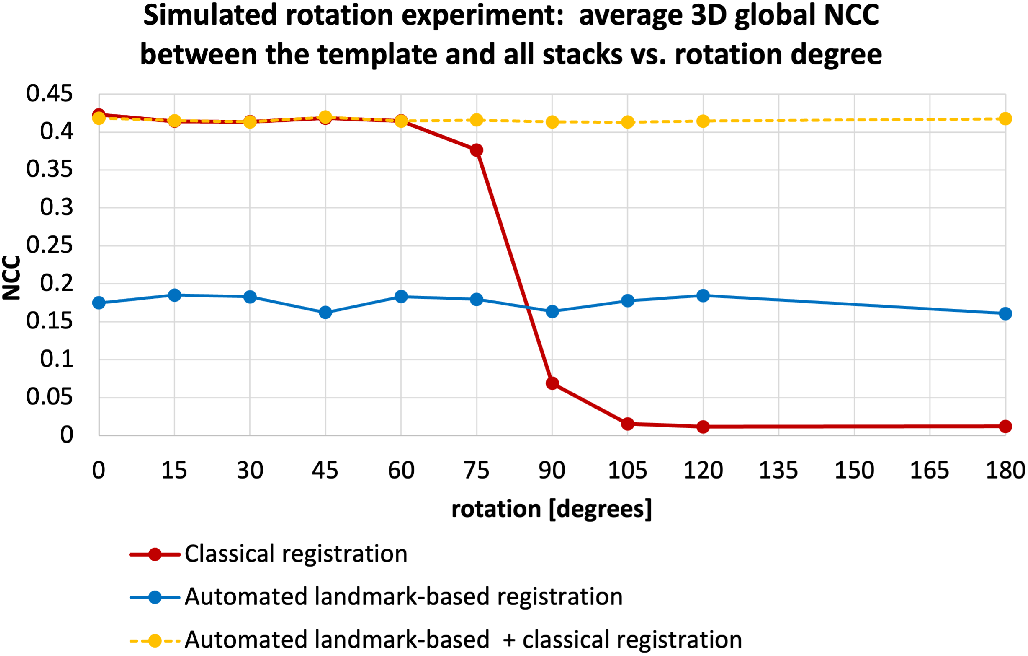
Simulated [0; 180] degrees range rotation experiment for comparison of the capture range of the classical rigid registration (red), automated landmark-based solution (blue) and combination of the classical registration initialised with the automated output (yellow): average global 3D NCC between the template and all transformed registered stacks in the masked thorax ROI.

Combination of these methods by initialisation of the classical regression-based registration with the global landmark-based pose estimation provides a stable high quality solution for the whole rotation range (yellow, NCC = 0.411 ± 0.083). At the 90 - 180 degree range, the landmark-based approach outperforms the classical rigid registration and the combined approach providing the best solution. The results are significant with *p* < 0.001. At the remaining part of the range there is no significant difference between the only classical and combined registration approaches.

Therefore, in the proposed full reconstruction pipeline, additional registration to the common space is used as a part of the stack selection step described in Sec. 2.4.

### 4.5. Automated DSVR reconstruction: severe motion datasets

The performance of the full proposed pipeline for automated reconstruction was evaluated on 5 cases from 21 - 24 weeks GA range and affected by severe motion with large rotations and translations. For these cases the original SVR-based reconstruction pipeline (Lloyd et al., 2019) failed (e.g., see Fig. 2) and the standard DSVR-based pipeline (Uus et al., 2020) led to exclusion of large (> 50%) proportion of slices that resulted in low reconstruction quality. Each of the datasets contains 6 stacks acquired under different orientations with respect to the uterus and the fetus. Prior to processing, all stacks were divided into four packages (groups of slices acquired consecutively and covering the whole ROI) based on the interleaved slice acquisition order.

We investigated 3 scenarios for automated reconstruction with different combinations of the pipeline components: (i) thorax localisation only with the least motion-corrupted stack selected as a template followed by DSVR (steps I+IV); (ii) reorientation of all stacks to the standard space with the least motion-corrupted stack selected as a template followed by DSVR (steps I+II+IV); (iii) full proposed pipeline (steps I+II+III+IV). In scenarios (i) and (ii), the template stack was automatically selected based only on the degree of motion corruption (NCC between sequential slices) similarly to the approach proposed in (Kainz et al., 2015). The results were quantitatively evaluated in terms of the total proportion of excluded slices, average global 2D NCC between all simulated and original slices and average 2D NCC between the simulated and original slices of an excluded stack (similarly to the approach in Uus et al. (2020)). The excluded stack was selected so that the thorax would be in approximately the same position as in the template to make sure that the main structures will be preserved and that it would not be completely misaligned. We selected NCC between the simulated and original slices for evaluation because registration in DSVR pipeline is driven by normalised mutual information metric.

The corresponding results presented in Tab. 2 show that addition of reorientation to the standard space (Step II) to the to localisation-only scenario (Step I) decreases the proportion of the excluded slices which depends on the quality of registration and increases the average NCC between for all slices in the dataset. It should be noted that the structure-based outlier rejection in the standard DSVR method was already designed for exclusion of the majority of misregistered slices and the localisation-only pipeline (Step I) is operational even in the extreme motion cases.

**Table 2.** Evaluation of the components of the proposed automated DSVR reconstruction pipeline on 5 severe rotation and translation motion datasets with respect to the % proportion of rejected slices and 2D NCC between simulated and original slices in all stacks and only the excluded stack. The investigated scenarios include: (i) Steps I+IV; (ii) Steps I+II+IV; (iii) full pipeline: Steps I+II+III+IV. The results are statistically significant with *p* < 0.001 for comparison between (i) vs. (ii) and (iii) scenarios and with *p* < 0.01 for (ii) vs. (iii) comparison. Step I is thorax localisation, Step II is reorientation to the atlas, Step III is stack selection and average template generation and Step IV is DSVR reconstruction.

| Scenario | % of rejected | all NCC | excl. NCC |
| --- | --- | --- | --- |
| I+IV | $64.7 \pm 5.3 \%$ | $0.579 \pm 0.058$ | $0.550 \pm 0.218$ |
| I+II+IV | $51.6 \pm 8.0 \%$ | $0.728 \pm 0.090$ | $0.705 \pm 0.105$ |
| I+II+III+IV | $37.8 \pm 9.1 \%$ | $0.834 \pm 0.028$ | $0.809 \pm 0.058$ |

The main causes of excluded slices are related to the intensity artefacts due to motion as well as misregistrations. Complete exclusion of outliers ensures that the intensity and registration errors are not propagated into the reconstructed volumes. But this also leads to the loss of information content required for super-resolution reconstruction (lower NCC in the excluded stack in scenarios I+IV and I+II+IV).

The full pipeline with the stack selection and template generation components (Step III) resulted in the highest NCC values between the simulated and original slices. This step refined global pose stack transformations (see. Sec. 4.4) while definition of the common average template space provided more stable initial registration target which led to the higher number of included slices.

An illustration of reconstruction results for one of the early GA datasets (23 weeks) affected by severe > 90 degrees rotation and translation motion is presented in Fig. 15. In comparison to the failed output (A) of the classical rigid SVR pipeline (Lloyd et al., 2019; Kuklisova-Murgasova et al., 2012), even without reorientation, the automated DSVR pipeline could still reconstruct the main anatomical features (B). However, this led to rejection of a large proportion of slices (58.8%) and grainy unstable texture, which made interpretation challenging. Addition of the reorientation step (C) reduced the number of excluded slices (38.7%) but the template was not optimally selected resulting in a blurred image. Finally, the proposed step for selection of stacks and generation of the average template (D) improved definition of the fine vascular structures due to the higher number of included slices (only 30.8% were excluded).

**Fig. 15.**
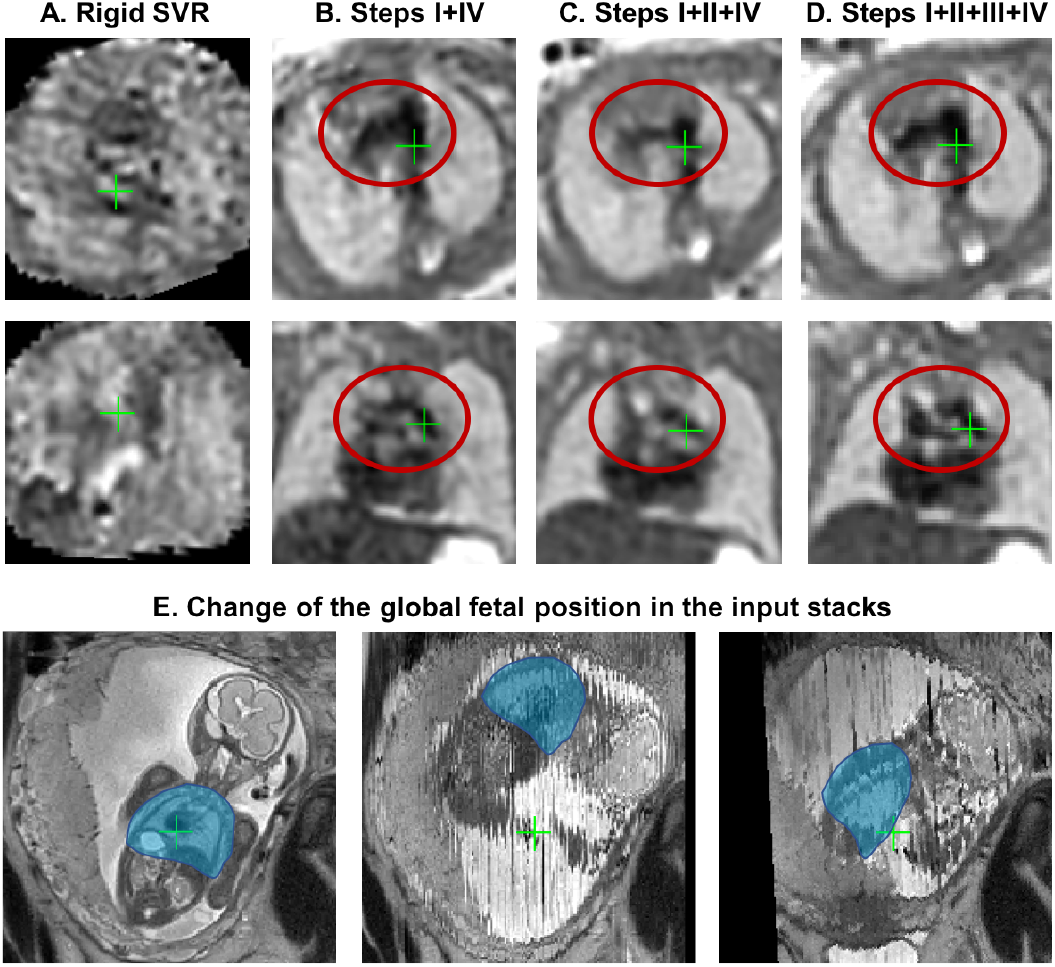
An example of reconstruction results for an early GA (23 weeks) dataset with 6 stacks affected by > 90 degrees rotation motion: (A) original manual rigid SVR pipeline (Lloyd et al., 2019), (B) Steps I+IV, (C) Steps I+II+IV, (D) full pipeline with Steps I+II+III+IV. Note that all images were additionally aligned to the same space for visualisation purposes (axial and coronal views). The global change of the fetal thorax (blue) position between the different input stacks in this dataset is shown in (E).

### 4.6. Qualitative evaluation: early and late GA datasets

The performance of the proposed pipeline was also qualitatively evaluated on 50 early GA datasets (≤ 24 weeks) from healthy controls of the iFIND project and 50 late GA datasets (≥ 30 weeks) from the clinical CMR practice where research consent was obtained.

The early GA-specific cohort is particularly prone to large rotations and translations that cannot be resolved by the classical registration methods, which effectively limited the previous 3D SVR fetal cardiac MRI study to primarily late GA (median ~ 32 weeks) cases (Lloyd et al., 2019). Furthermore, one of the conditions for a stable reconstruction of small vascular structures in this subject group is the inclusion of all available image information and minimisation of the proportion of excluded slices.

The selection criteria out of all available iFIND datasets were: ≤ 24 weeks GA, singleton pregnancies, similar acquisition protocol (Sec.4.1), more than 5 available stacks, and no extreme SNR loss. The 50 late GA CHD CMR datasets (≥ 30 weeks) were selected randomly from the recent acquisitions with the consent for research and no extreme SNR loss. The reconstructions were performed using the full version of the proposed automated pipeline with 0.7mm output isotropic resolution. The output 3D volumes were graded by a clinician trained in fetal CMR in terms of both general image quality and visibility of the major cardiovascular structures essential for diagnosis of a specific group of CHD (major vascular abnormalities). The quality grading scheme (Fig. 16) has four categories {0; 1; 2; 3} with 0 corresponding to failed reconstruction, and 1, 2 and 3 to poor, good/adequate and high image quality, correspondingly. The datasets graded ≥ 2 are considered to be acceptable for detailed clinical assessment and interpretation of the specific group of CHD (major vascular abnormalities) with all major cardiovascular structures being clearly visible.

**Fig. 16.**
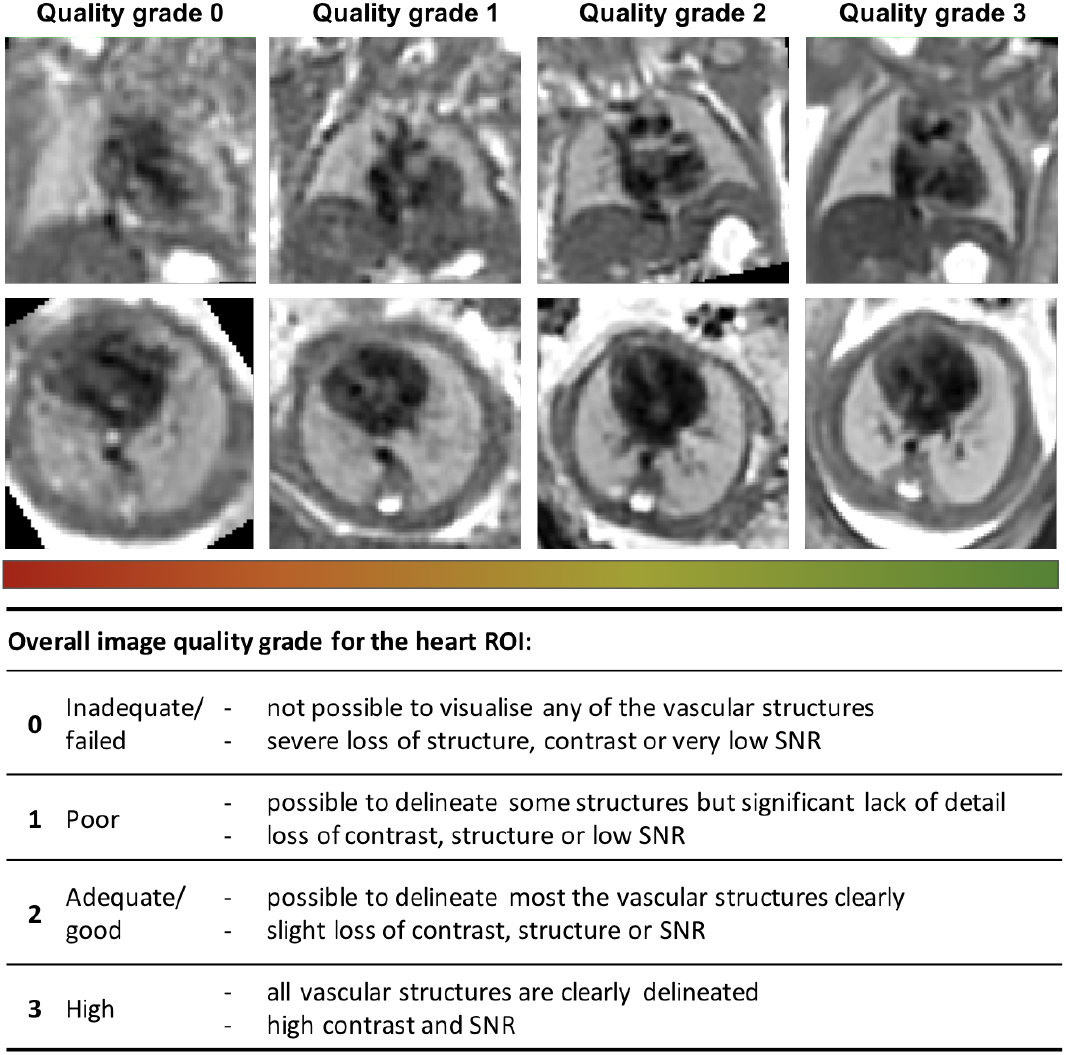
Fetal thorax reconstruction quality grading scheme for the heart ROI based on the proposed fully automated DSVR pipeline along with the examples from the early GA cohort.

The histogram of the quality grades for the early GA cohort presented in Fig. 17.B shows that the majority of the grades are within 2-3 range and therefore acceptable for interpretation, with the average grade 2.16 ± 0.68. The primary causes of the low grades in 6 cases are related to significantly lower SNR levels in the input stacks (low quality acquisition) in combination with the small vessel size.

The histogram of the reconstruction image quality grades for the late GA cohort is presented in Fig. 17.C and nearly all grades (apart from 1 case) are within 2-3 range and therefore acceptable for interpretation with the average grade 2.60 ± 0.53. As expected, the proportion of the cases with the high image quality score is higher than in the early GA cohort since the size of the vessels in significantly larger with respect to the reconstruction resolution.

The distribution of the quality scores of the manual DSVR reconstruction for the same early GA cohort is given in Fig. 18A. It should be noted that, in this experiment, we only created the manual masks and did not perform careful template selection (like it was done in (Uus et al., 2020)) but only chose the stacks from the middle of the dataset. The automated method improved the image quality in 36% of the cases and did not decrease the quality in any of the rest. The primary reasons for the lower manual reconstruction results include the fact that the template was chosen randomly (unlike the average in the proposed solution) and the impact of large rotations. This led to the lower degree of oversampling resulting the lower visibility of the vessels or even a complete failure of reconstruction (e.g., Fig. 18)B

**Fig. 17.**
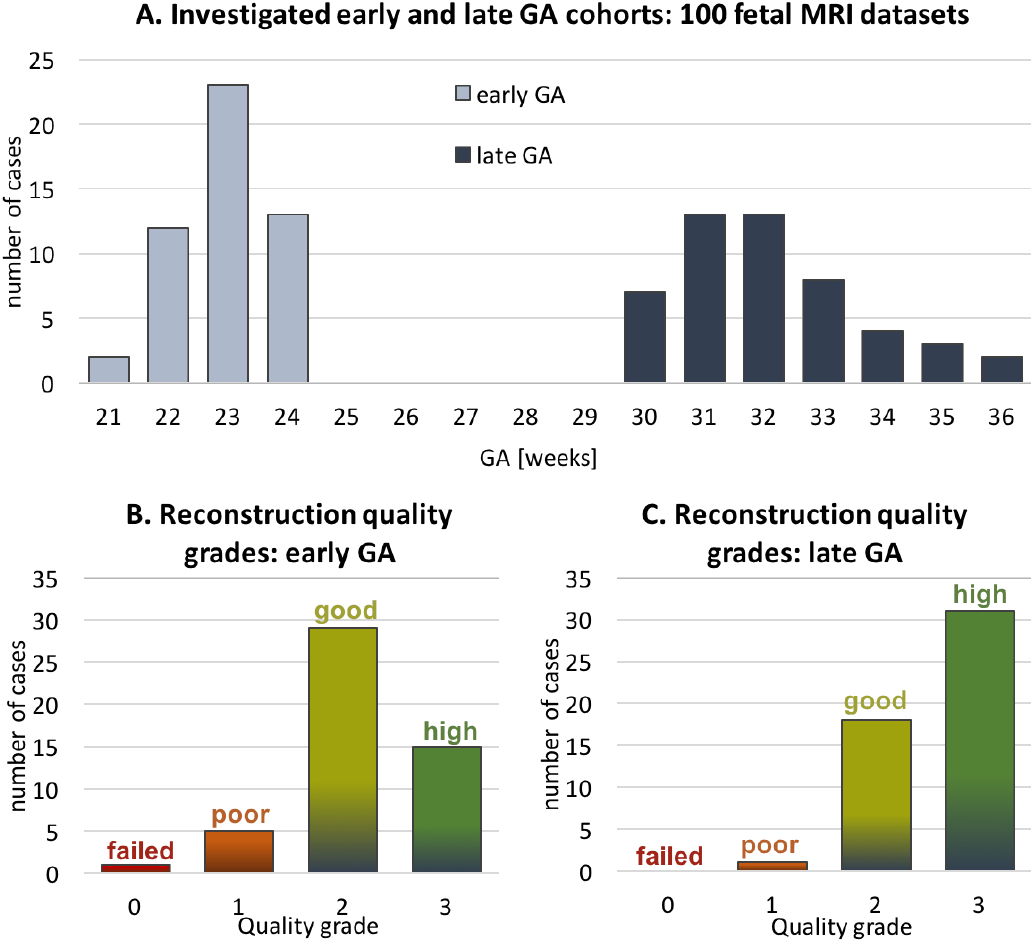
The results of qualitative assessment of fetal thorax reconstruction using the proposed automated DSVR pipeline: distribution of image quality scores for 50 early (B) 50 late GA (C) fetal MRI datasets. The GA distribution of the investigated datasets is presented in (A).

**Fig. 18.**
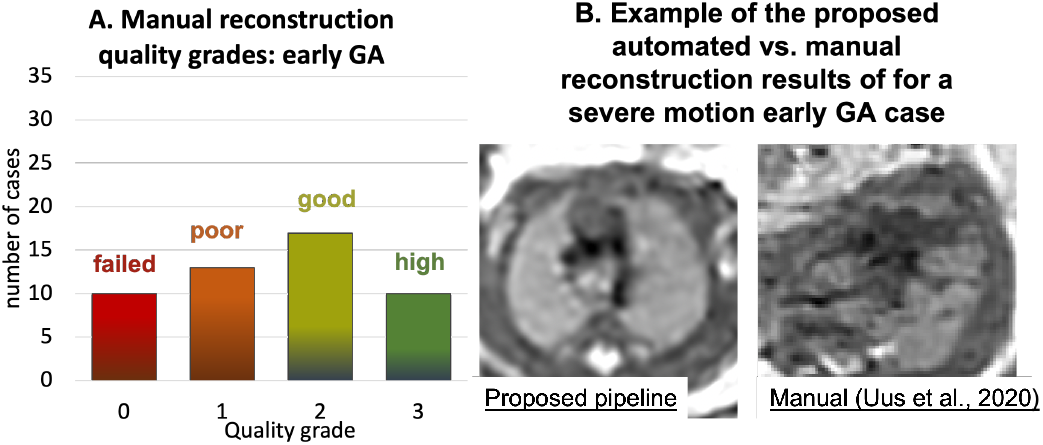
The results of qualitative assessment of fetal thorax reconstruction using the original manual DSVR for the early GA cohort (50 cases): distribution of image quality scores (A) and an example of manual vs. automated reconstruction with different grades (B).

In addition, Fig. 19 shows comparison of the outputs of the proposed automated DSVR-based pipeline with the classical manual rigid SVR reconstruction used in (Lloyd et al., 2019) for a clinical study on two 23 weeks GA cases affected by severe and minor rotations. The visual assessment of the results shows the superior image quality and sharper features even for the minor motion dataset.

**Fig. 19.**
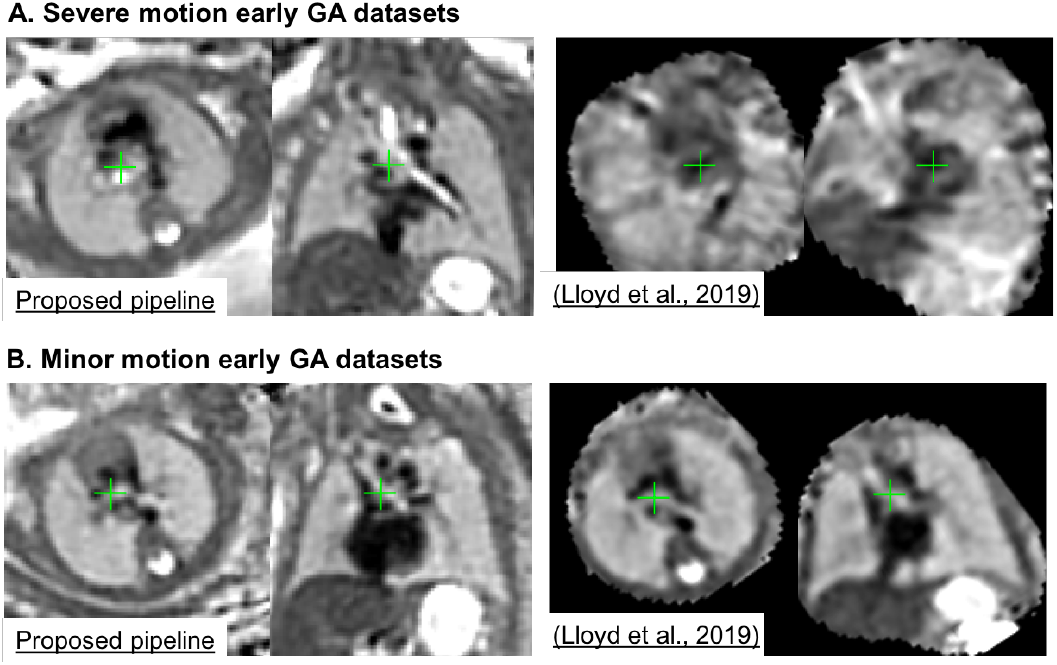
An example of the proposed automated DSVR pipeline vs. the classical manual rigid SVR reconstruction (used for clinical studies in (Lloyd et al., 2019)) for severe (A) and minor (B) rotation and translation motion early GA (23 weeks) datasets.

The selected set of landmarks (Sec.2.3) is also operational for the cases with pronounced anomalies that significantly change the anatomy. Although it might required additional reorientation to the atlas space due to the shifted landmark positions. For example, Supplemental Figure S3 demonstrates reconstruction results for congenital diaphragmatic hernia. While the shifted relative liver and heart positions led to slightly different orientation of the output image the reconstruction quality is high.

## 5. Discussion and limitations

Automation of 3D (D)SVR-based reconstruction process and correction of extreme motion are reportedly the two major challenges in fetal MRI motion correction. And while the existing CNN-based solutions for the fetal brain (e.g., (Salehi et al., 2019)) already showed promising results, these methods have not yet been applied to the fetal trunk ROI.

This work focused on development of a practical solution for automated DSVR reconstruction of the fetal thorax combining 3D CNN-based intra-uterine localisation of the fetal trunk in motion corrupted stack followed by automated reorientation of the fetal thorax to the standard atlas space using 3D CNN segmentation of a set or ROI-specific landmarks within the fetal trunk. The reoriented stacks are then passed to the classical DSVR reconstruction (Uus et al., 2020) with an additional automated stack selection and template generation steps based on motion corruption and mutual stack similarity metrics.

We demonstrated that the proposed localisation pipeline based on the multi-label 3D UNet provides robust 3D detection of the trunk even in stacks with partial fetal coverage. The landmark-based solution is also efficient for global fetal pose estimation and extends the rotation and translation capture range of the classical rigid registration. The proposed step for automated generation of the template space and exclusion of potentially misregistered or low image quality stacks also showed to improve reconstruction quality.

In addition, the pipeline was qualitatively evaluated on 100 randomly selected fetal MRI datasets from 21 to 36 week GA range in terms of the image quality acceptable for anatomical interpretation of the major cardiovascular structures. The results showed that the majority of the early GA datasets (88%) have good image quality with clearly defined most of the major cardiovascular structures. The poor reconstruction quality in the remaining proportion of cases is related to the low input image quality in combination with the small vessel size at this age range which emphasises the need for optimisation of the super-resolution reconstruction step and correction of intensity artefacts. The automated DSVR-based pipeline also produced superior image quality in comparison to the conventional manual rigid SVR-based approach even for minor motion cases. In addition, the performed assessment of 50 late GA CMR cases showed high image reconstruction quality with well defined vascular structures. These results indicate the potential feasibility to extend the application of 3D fetal SVR/DSVR reconstruction-based CMR analysis to the wider GA range which is currently primarily limited to the late GA cases (Lloyd et al., 2019) due to the extreme rotation motion affecting young fetus datasets.

In terms of the limitations of the proposed solution, it should be noted that the image quality and degree of motion corruption directly affect segmentation and landmark estimation accuracy with potential errors propagating to the registration step. Therefore, the landmark-based approach is limited by the condition that the individual fetal trunk structures should be identifiable in all stacks. Comparison to the more traditional regression-based fetal pose estimation methods (e.g., (Hou et al., 2018)) should also be investigated since it might provide an alternative solution for severely motion-corrupted stacks. This is planned to be addressed in our future work.

Other aspects such as low image quality (low SNR or intensity artefacts) and different acquisition protocols would require further training of the networks on a wider range of datasets with different acquisition parameters and range of anomalies.

Although, both of the aforementioned limitations (errors in registration or low image quality) can be resolved by rejection of outliers this would still lead to loss to useful information required for reconstruction of fine vascular structures. In order to minimise this loss of information, an optimal solution should include advanced signal processing methods for reconstruction of the datasets affected by low SNR or severe intensity artefacts.

Quantitative evaluation of the per-vessel image quality (in addition to the qualitative study in Section 4.6) is a challenging step because there is no ground truth. It can be assessed only with respect to visibility of individual extracardiac vascular structures (used in diagnosis in the current fetal 3D T2w CMR protocol). The presented qualitative evaluation score reflects the opinion of the clinical experts in 3D fetal heart MRI (Lloyd et al., 2019).

## 6. Conclusions

In this work, we proposed and implemented a first fully automated pipeline for robust DSVR reconstruction of high resolution 3D fetal thorax images from motion-corrupted T2-weighted MRI stacks. It based on CNN-based solutions for automated localisation and pose estimation for correction of large magnitude motion for the fetal trunk ROI, which was not achievable before. Furthermore, the reconstruction process is performed directly in the standard atlas space. The pipeline was quantitatively evaluated on a series of fetal MRI datasets and a simulated experiment. We also performed qualitative assessment on 100 early and late GA fetal MRI datasets with the image quality grading results suggesting the potential feasibility of using 3D automated DSVR reconstructions for clinical interpretation of the major cardiovascular structures.

## Acknowledgments

We thank everyone who was involved in acquisition and examination of the datasets and all participating mothers.

This work was supported by the Rosetrees Trust [A2725], the Wellcome/EPSRC Centre for Medical Engineering at King’s College London [WT 203148/Z/16/Z], the Wellcome Trust and EPSRC IEH award [102431] for the iFIND project, the NIHR Clinical Research Facility (CRF) at Guy’s and St Thomas’ and by the National Institute for Health Research Biomedical Research Centre based at Guy’s and St Thomas’ NHS Foundation Trust and King’s College London.

The views expressed are those of the authors and not necessarily those of the NHS, the NIHR or the Department of Health

## Footnotes

2 PyTorch: https://pytorch.org

3 SVRTK fetal MRI segmentation repository: https://github.com/SVRTK/Segmentation_FetalMRI

4 MIRTK library: https://github.com/BioMedIA/MIRTK

5 SVRTK toolbox: https://github.com/SVRTK/SVRTK

6 SVRTK fetal thorax reconstruction docker (*fetal_thorax* tag): https://hub.docker.com/repository/docker/fetalsvrtk/svrtk

7 iFIND project: https://www.ifindproject.com

