## Supplemental materials for "Automated 3D reconstruction of the fetal thorax in the standard atlas space from motion-corrupted MRI stacks for 21-36 weeks GA range"

### Supplemental material

**Figure S1.** An example of comparison of single label fetal trunk localisation results for 2D vs. 3D UNet (an earlier experiment). The segmentation outputs are visualised as red. The results are statistically significant with $p < 0.0001$.

**Comparison of 2D vs. 3D UNet for fetal trunk localization in fetl MRI stacks**

- **Training was performed on:** 30 datasets with 257 stacks with TorchIO augmentation
- **Testing was performed on:** 20 stacks covering the fetal trunk ROI

| Metric | 2D vs. GT | 3D vs. GT |
|---|---|---|
| Dice | 0.463 ± 0.100 | 0.863 ± 0.061 |
| Sensitivity | 0.743 ± 0.146 | 0.937 ± 0.040 |
| Specificity | 0.968 ± 0.009 | 0.995 ± 0.040 |

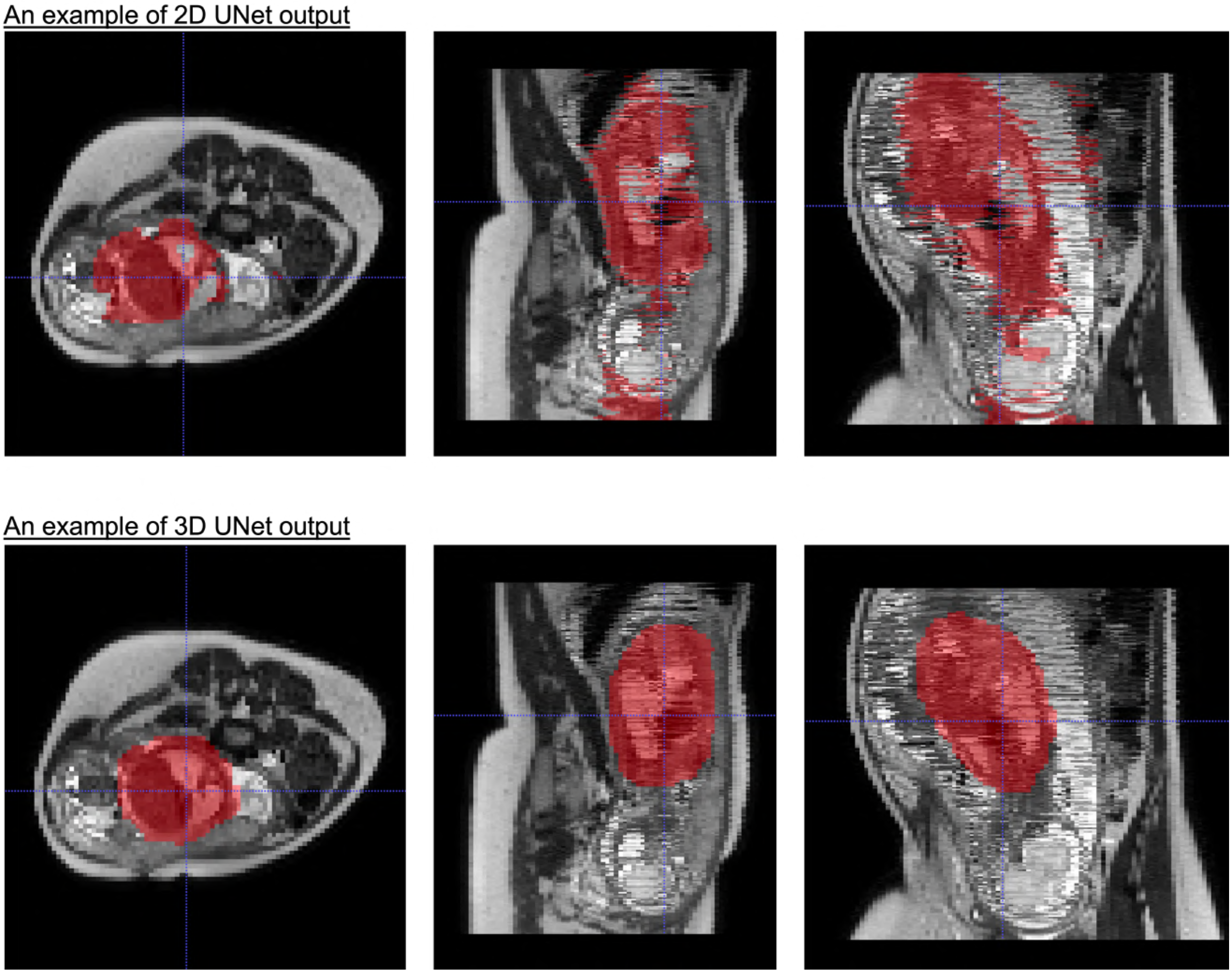

**Figure S2.** An example of the impact of the number of stacks on the output image quality for a minor motion dataset. The reconstruction quality improves with higher number of stacks due to oversampling of the region of interest.

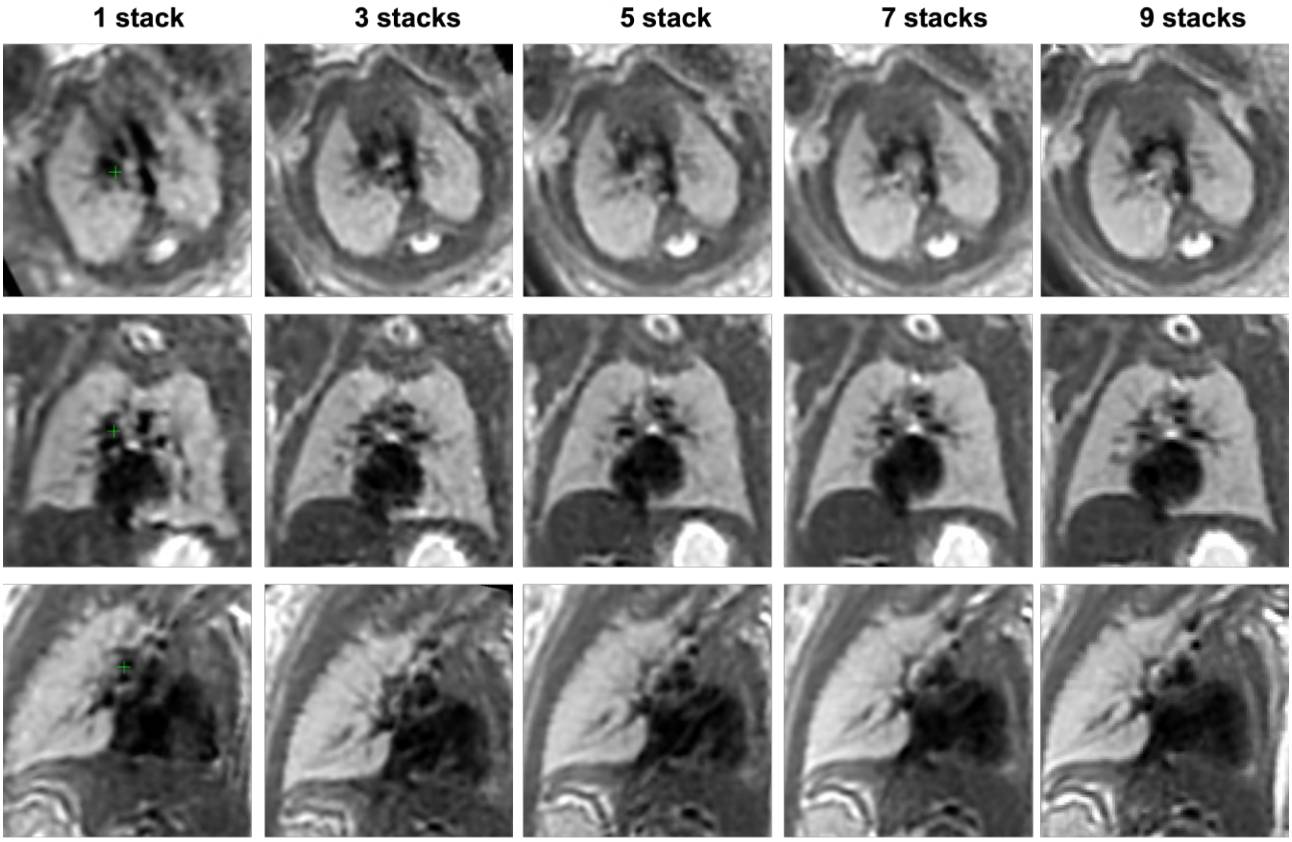

**Table ST1.** Description of the datasets used for each of the experiments in Sections 4.2-6 including both training of the networks and testing.

| Purpose | Input data format | Training/Validation | Testing |
|---|---|---|---|
| **Training and localisation network (1) for global fetal position: uterus, trunk and brain** | - Raw CMR and iFIND datasets: early and late GA<br>- Manual labels of uterus, trunk and head for each of the stacks | 35 CMR datasets with 354 stacks in total | Experiment 1 (Section 4.2): 16 CMR and iFIND datasets with 164 stacks in total |
| **Localisation networks (2) and 3D for local landmarks: hear, liver, thorax and abdomen** | - DSVR reconstructed images (iFIND) and raw MRI stacks (CMR) cropped to the trunk ROI: early and late GA<br>- Landmark labels (thorax, abdomen, heart, liver) propagated from the atlas (and refined, if required) | 70 cases from (Uus et al., 2020) | Experiment 2 (Section 4.3): 50 raw iFIND stacks from different datasets |
| **Simulated experiment for assessment of the rotation capture range** | Raw stacks cropped to the thorax ROI: late GA | -------------------- | Experiment 3 (Section 4.4): 5 datasets with 7 stacks each |
| **Quantitative assessment of required quality for severe motion early GA cases** | Raw iFIND datasets: early GA | -------------------- | Experiment 4 (Section 4.5): 5 datasets with 6 stacks each |
| **Qualitative assessment of reconstruction quality** | Raw CMR and iFIND datasets : early and late GA | -------------------- | Experiment 5 (Section 4.6): 100 CMR and iFIND datasets |

**Figure S3.** An example of the automated reconstruction results for congenial diaphragmatic hernia dataset. While the shifted relative liver and heart positions (A) led to slightly different orientation of the output image the reconstruction quality is high (B). The rotation can be resolved by additional reorientation to the true atlas space (C).

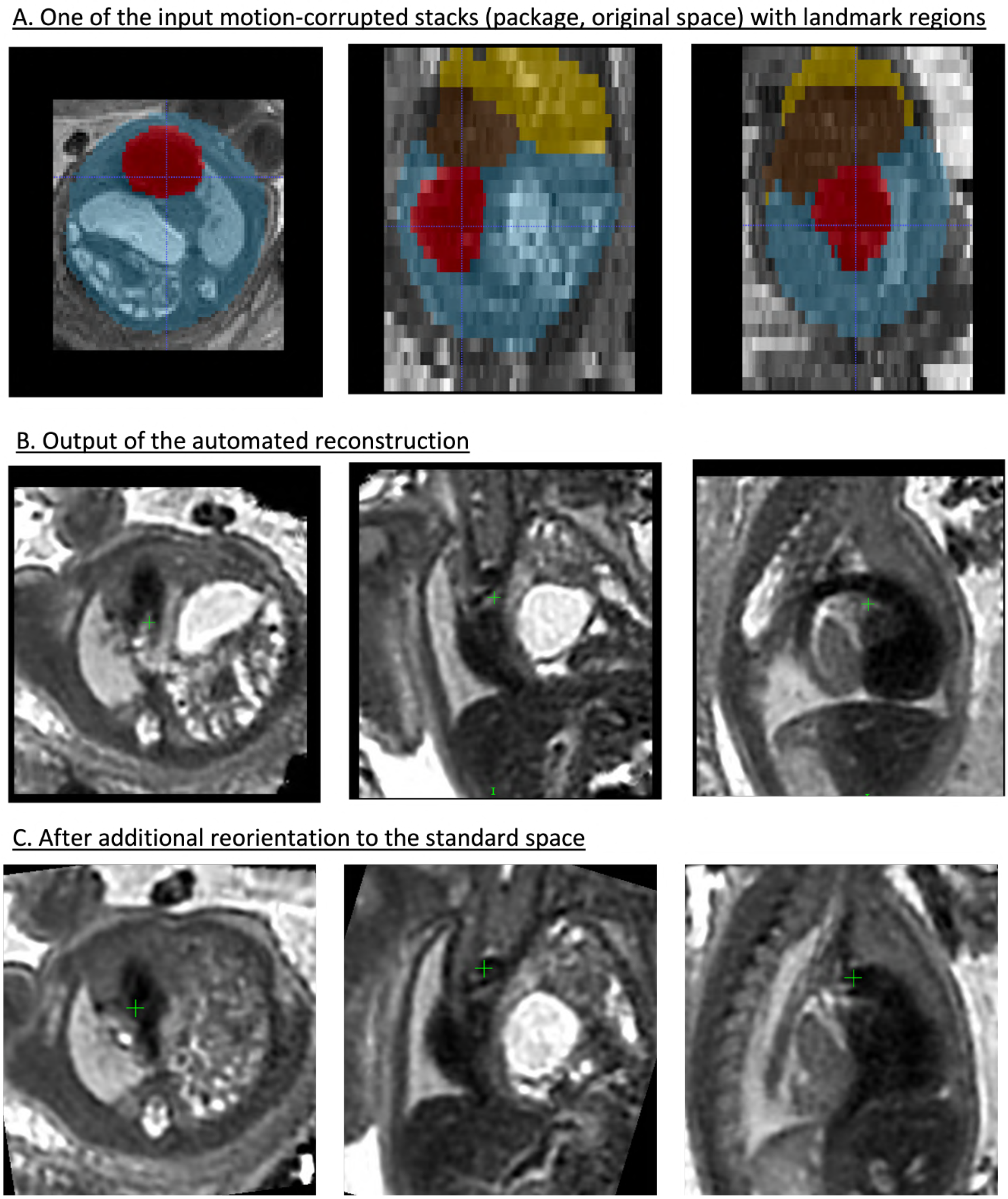
